# The functional evolution of termite gut microbiota

**DOI:** 10.1101/2021.12.01.470864

**Authors:** Jigyasa Arora, Yukihiro Kinjo, Jan Šobotník, Aleš Buček, Crystal Clitheroe, Petr Stiblik, Yves Roisin, Lucia Žifčáková, Yung Chul Park, Ki Yoon Kim, David Sillam-Dussès, Vincent Hervé, Nathan Lo, Gaku Tokuda, Andreas Brune, Thomas Bourguignon

## Abstract

Termites primarily feed on lignocellulose or soil in association with specific gut microbes. The functioning of the termite gut microbiota is partly understood in a handful of wood-feeding pest species, but remains largely unknown in other taxa. We intend to feel this gap and provide a global understanding of the functional evolution of termite gut microbiota. We sequenced the gut metagenomes of 145 samples representative of the termite diversity. We show that the prokaryotic fraction of the gut microbiota of all termites possesses similar genes for carbohydrate and nitrogen metabolisms, in proportions varying with termite phylogenetic position and diet. The presence of a conserved set of gut prokaryotic genes implies that key nutritional functions were present in the ancestor of modern termites. Furthermore, the abundance of these genes largely correlated with the host phylogeny. Finally, we found that the adaptation to a diet of soil by some termite lineages was accompanied by a change in the stoichiometry of genes involved in important nutritional functions rather than by the acquisition of new genes and pathways. Our results reveal that the composition and function of termite gut prokaryotic communities have been remarkably conserved since termites first appeared ∼150 million years ago. Therefore, the “world smallest bioreactor” has been operating as a multipartite symbiosis composed of termites, archaea, bacteria, and cellulolytic flagellates since its inception.

## INTRODUCTION

Termites are one of the few animal lineages feeding on substrates distributed along the wood-soil decomposition gradient (Donovan *et al*., 2001; Bourguignon *et al*., 2011). Although termites produce their own endogenous cellulases (Watanabe *et al*., 1998; Tokuda *et al*., 2004), their ability to decompose wood or soil organic matter largely depends on symbiosis with mutualistic gut microbes (Watanabe and Tokuda, 2010; Brune and Ohkuma, 2011), including bacteria, archaea and, in the case of lower termites, cellulolytic flagellates. The cellulolytic flagellates of termites are typically found nowhere else other than in termite guts and are efficiently transmitted across host generations (Nalepa, 2017; Michaud *et al*., 2020). Similarly, many of the prokaryotes present in termite guts are found nowhere else in nature (Bourguignon *et al*., 2018; Hervé *et al*., 2020). Their vertical mode of inheritance is supported by the observations that differences among termite gut prokaryotic and protist communities tend to increase as phylogenetic distances among termite hosts increase (Rahman *et al*., 2015; Tai *et al*., 2015). In addition, the diet of the termite host, which largely correlates with the termite phylogeny (Bourguignon *et al*., 2011), also shapes the termite gut microbial communities (Dietrich *et al*., 2014; Mikaelyan *et al*., 2015). Whether the termite phylogeny is recapitulated by gut microbial functions, as it is recapitulated by the taxonomic composition of microbial communities, remains unknown.

Investigations of termite gut microbe genomes has revealed that, in addition to the production of enzymes involved in lignocellulose digestion, gut microbes have numerous nutritional functions, including nitrogen fixation and recycling abilities that supplement the nitrogen-poor diet of their host (Lilburn *et al*., 2001; Yamada *et al*., 2007; Hongoh *et al*., 2008; Ohkuma, M. & Brune, 2011). While metagenomics and metatranscriptomics surveys of termite guts have been carried out for an increasingly large number of termite species (Warnecke *et al*., 2007; He *et al*., 2013; Liu *et al*., 2018; Tokuda *et al*., 2018; Marynowska *et al*., 2020), often with the prospect of harvesting cellulolytic enzymes able to convert plant biomass into biofuel (*e.g.* Tartar *et al*., 2009; Calusinska *et al*., 2020), there has been a marked sampling bias towards easy-to-sample wood-feeding termite species, and species with pest status. Far less is known about the function and taxonomy of the gut prokaryotic communities of other termite lineages, such as basal wood-feeding lineages, or lineages with soil-feeding habits (Hervé *et al*., 2020). Because of this gap in our knowledge, it remains largely unclear how the taxonomy and function of gut microbiome has been evolving since termites came to be ∼150 Million years ago (Bourguignon *et al*., 2015; Bucek *et al*., 2019). Similarly, how the acquisition of a diet based on soil has affected the taxonomy and function of gut microbial communities remains an open question. A metagenomics survey based on a comprehensive sampling of termites is required to answer these questions.

In this study, we sequenced whole gut metagenomes of 145 termite samples representatives of the phylogenetic and ecological diversity of termites, including many lineages that have remained undocumented. We also sequenced the gut metagenome of one sample of *Cryptocercus*, the sister group of termites (Lo *et al*., 2000). We used the assembled prokaryotic contigs of this dataset to determine (1) when important gut prokaryotic pathways involved in nutritional functions were acquired by termites; (2) to which extent termite phylogeny is predictive of gut prokaryote taxonomic and functional composition; and (3) the taxonomic and functional changes experienced by gut prokaryote communities following the acquisition of a diet of soil.

## RESULTS AND DISCUSSION

### The taxonomic composition of termite gut prokaryotes

We sequenced whole gut metagenomes, including the hindgut containing the bulk of the gut microbiota, of 145 termite species (Table S1, Figure S1). This included species from the nine termite families and species from the eight subfamilies of Termitidae (Lo *et al*., 2000). Our shotgun sequencing approach generated an average of 72.5 million reads per sample that were assembled into an average of 92,237 scaffolds >1000 bps, constituting 63.3% of mapped reads. The proportions of prokaryotic reads were on average 18.4% in lower termites and 20.5% in higher termites.

We used 40 marker genes (Sunagawa *et al*., 2013; Wu *et al*., 2013) to determine the taxonomy and estimate the abundance of each major bacterial lineage present in the 129 termite gut metagenome assemblies including upward of 10,000 contigs longer than 1000 bps. Shorter contigs were removed from the analyses. The bacterial community composition and abundance inferred from marker gene data showed similarities at the phylum level to that inferred from 16S rRNA gene amplicon sequences (Figure S2). However, the abundance distribution estimated by both approaches showed some disagreements for several families (Dietrich *et al*., 2014; Mikaelyan *et al*., 2015; Bourguignon *et al*., 2018). Notably, *Dysgomonadaceae*, *Ruminococcaceae*, *Synergistaceae*, and *Oscillospiraceae* occurred at low abundances among the marker genes but were represented by many 16S rRNA gene sequences in most termite species (Dietrich *et al*., 2014; Mikaelyan *et al*., 2015; Bourguignon *et al*., 2018) (Table S2). These discrepancies are likely the result of variation in 16S rRNA gene copy number (Větrovský and Baldrian, 2013; Edgar, 2018), which are higher in these lineages, or are possibly artifacts generated during 16S rRNA gene amplicon PCR cycles. They might also reflect the incomplete coverage of our metagenomes or, to a certain extent, the differences in the databases used for classification.

In total, we identified 114 family-level bacterial lineages, belonging to 19 phyla and represented in the gut of more than 5% of termite species (Table S3). Many other bacterial family-level lineages were recorded from the gut of no more than a few termite species, and were possibly transient, and not strictly associated with termite guts. We calculated the Moran I index on the abundance of these 114 family-level bacterial lineages to test whether bacterial abundance is correlated with termite phylogeny. We found a phylogenetic autocorrelation signal for 59 of the 114 bacterial lineages, and this signal remained significant at a 5% false discovery rate (FDR) correction for 27 bacterial lineages, including some of the most abundant bacterial lineages (Figure 1, Table S4). For example, the wood-fiber-associated *Fibrobacteraceae* (Mikaelyan *et al*., 2014; Tokuda *et al*., 2018) are dominant in the gut of *Microcerotermes*, Nasutitermitinae, and related termite lineages, and are either undetectable or occur at low abundance in the assemblies of other termite lineages. Another example is the *Endomicrobiaceae* that comprise flagellate-associated (Stingl *et al*., 2005; Zheng *et al*., 2015) and free-living Endomicrobia (Ikeda-Ohtsubo *et al*., 2016; Mikaelyan *et al*., 2017), which were abundant in lower termites and almost entirely absent in higher termites.

**Figure 1.**
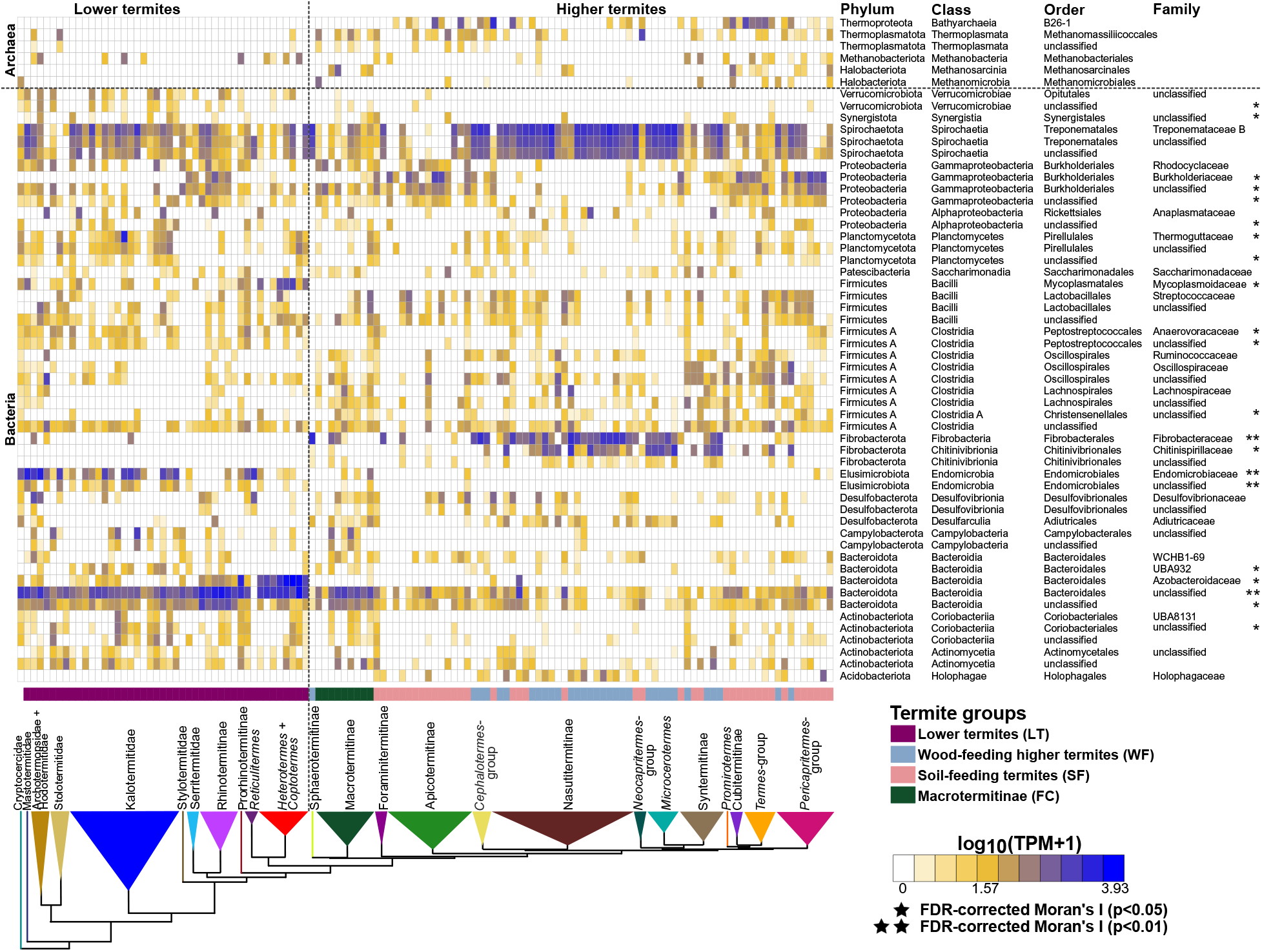
Relative abundance of the top 50 bacterial lineages and the major archaeal orders found in the gut metagenomes of termites. The relative abundance of prokaryotic taxa was inferred from 40 single-copy marker genes. The color scale represents the logarithm of transcripts per million (TPM). The tree represents a simplified time-calibrated phylogenetic tree reconstructed using host termite mitochondrial genome sequences. Prokaryotic taxa presenting significant phylogenetic autocorrelation with the host phylogeny at a 5% false discovery rate (FDR) are indicated with an asterisk (*p < 0.05; **p < 0.01).

Our dense taxonomic sampling of diverse termite hosts also allowed us to identify bacterial lineages whose association with termites has remained largely unreported. For example, we found that the *Holophagaceae*, a bacterial family of Acidobacteriota previously reported from the gut of three humus-feeding termite species (Mikaelyan *et al*., 2015) and two species of Nasutitermitinae (Dietrich *et al*., 2014), is widely distributed in Nasutitermitinae, Foraminitermitinae, the *Cephalotermes*-group, and the *Pericapritermes*-group (Figure 1). Altogether, our results demonstrate that termite phylogeny is remarkably predictive of the gut bacterial community composition, as has been demonstrated for termite gut protists (Tai *et al*., 2015).

Using the same 40 marker genes and 129 metagenome assemblies used for bacteria, we investigated the diversity of gut-associated Archaea across the termite phylogenetic tree. In total, we identified 16 family-level archaeal lineages, including *Methanoculleaceae* and *Methanocorpusculaceae* (order *Methanomicrobiales*), *Methanosarcinaceae* (order *Methanosarcinales*), *Methanobacteriaceae* (order *Methanobacteriales*), *Methanomethylophilaceae* (order *Methanomassiliicoccales*), and *UBA233* (class *Bathyarchaeia*). All but nine family-level lineages were present in the gut of more than 5% of termite species. The abundance of *Methanosarcinaceae*, *UBA233*, and an unclassified family-level lineage of *Bathyarchaeia* showed significant autocorrelation signals with the termite phylogenetic tree when no FDR correction was applied (Figure 1, Table S4). *Bathyarchaeia* occurred in the clade of Termitidae excluding Macrotermitinae, Sphaerotermitinae, and Foraminitermitinae confirming previous reports (Loh *et al*., 2021), and *Methanosarcinaceae* was found in Macrotermitinae, Nasutitermitinae, and in Cubitermitinae and related termite lineages (Figure 1). Archaea represented in average less than 1% of the gut prokaryotes in wood-feeding termite species, while their proportion reached 4.6% in Macrotermitinae and 10.6% in soil-feeding termite species, and was exceptionally high in the soil-feeding *Mimeutermes* in which 59.8% of the marker genes were assigned to *Bathyarchaeia*. Our results are in line with the higher archaeal-to-bacterial ratios reported in soil-feeding termites as compared to their wood-feeding counterpart, reflecting the higher methane emission rates of soil-feeding termites (Brune, 2018, 2019).

### The carbohydrate-active enzymes of termite gut prokaryotes

We investigated the evolution of prokaryotic carbohydrate-active enzymes (hereafter: CAZymes) using the same 129 gut metagenome assemblies used to investigate gut prokaryotic composition. The *de novo* assemblies of these 129 gut metagenomes contained an average of 127,159 prokaryotic open reading frames (ORF). We identified ORFs coding for CAZymes using Hidden Markov model searches against the dbCAN2 database (Zhang *et al*., 2018). As a first step, we investigated the evolution of enzymes derived from prokaryotes with no consideration of their taxonomic origin. In total, we found 346 CAZyme categories in 129 gut metagenomes that consisted of 205 glycoside hydrolases (GHs), 57 glycoside transferases (GTs), 18 enzymes with carbohydrate-binding modules (CBMs), 16 carbohydrate esterases (CEs), 41 polysaccharide lyases (PLs), and 9 redox enzymes with auxiliary activities (AAs) (Table S5). We did not find any CAZymes in only one gut metagenome (that of *Araujotermes parvellus*, at e-value cut-off below e-30). For the other 128 gut metagenomes, the number of CAZyme categories varied between 5 and 139 per gut metagenome. Five GH families, GH2, GH3, GH10, GH31, and GH77, were found in more than 85% of the termite species. 14 GHs, seven of which had putative lignocellulolytic activity, were found in 75 to 85% of the termite species. Therefore, glycoside hydrolases previously found to be abundant in the gut of particular termite species (*e.g.* Warnecke *et al*., 2007, Calusinska *et al*., 2020) are universally part of the gut enzymatic repertoire of termites.

We calculated the Moran I index on the abundance of 211 CAZymes, including 146 CAZyme families and 65 sub-families, present in more than 10% of termite species, and found an autocorrelation signal with the termite phylogenetic tree for 107 CAZymes. The autocorrelation signal remained significant after FDR correction for 77 CAZymes (Figure 2, Table S6). Therefore, as for gut prokaryotic composition, termite phylogeny is predictive of the CAZyme repertoire present in termite guts.

**Figure 2.**
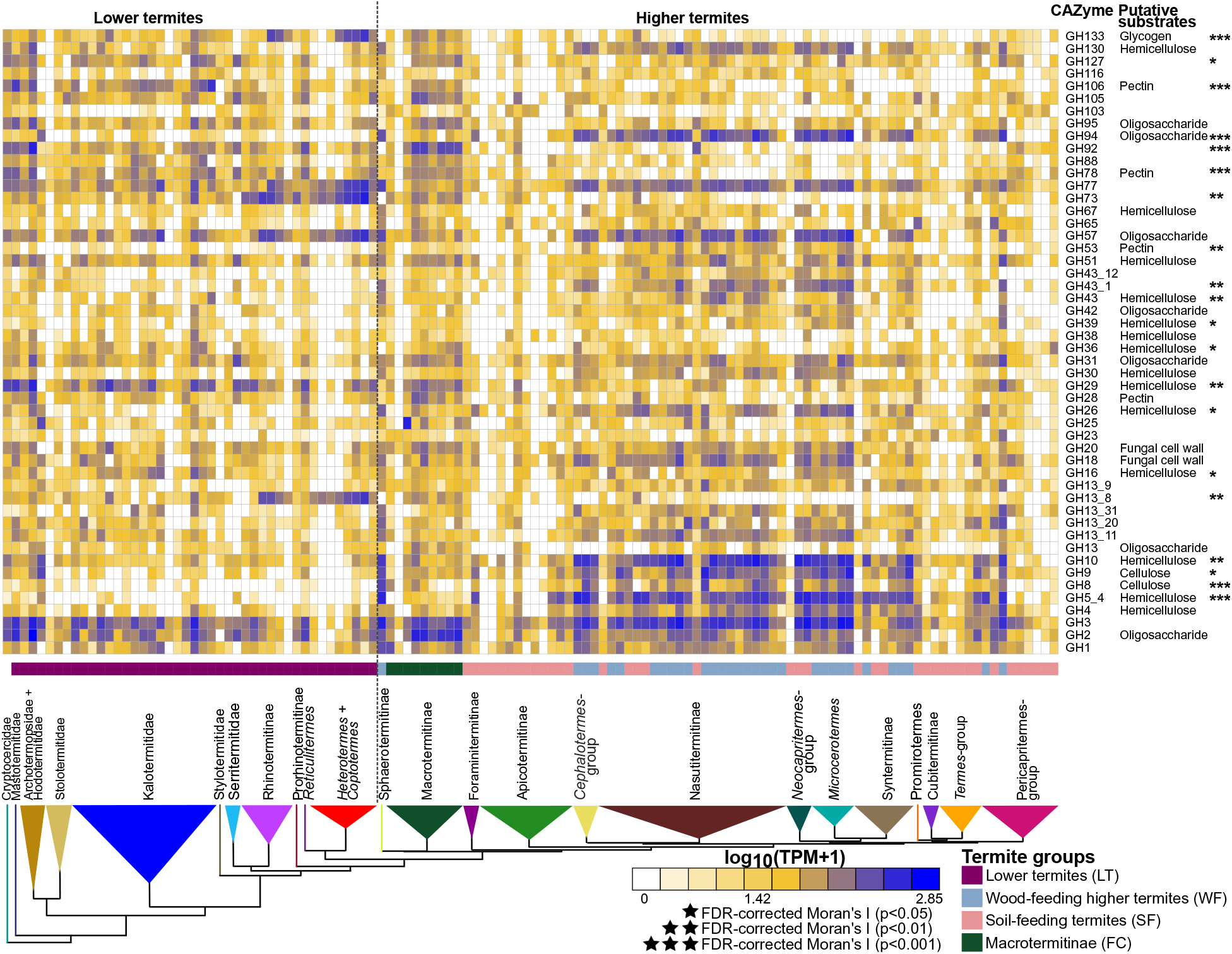
Relative abundance of CAZymes found in gut metagenomes of termites. The heatmap shows the 50 most abundant CAZymes. The color scale represents the logarithm of transcripts per million (TPM). The tree represents a simplified time-calibrated phylogenetic tree reconstructed using host termite mitochondrial genomes. Genes showing significant phylogenetic autocorrelation with the host phylogeny at a 5% false discovery rate (FDR) are indicated with asterisks (*p < 0.05; **p < 0.01).

Two factors that potentially affect the prokaryotic CAZyme repertoire of termite gut prokaryotes are diet and co-occurring non-prokaryotic cellulolytic symbiotic partners. We distinguished four termite groups: soil-feeding Termitidae (SF) and wood-feeding Termitidae excluding Macrotermitinae (WF), which host no other symbionts than gut prokaryotes (Brune, 2014), the fungus-cultivating Macrotermitinae (FC), which feed on wood or plant litter and cultivate cellulolytic fungi of the genus *Termitomyces* (Rouland-Lefèvre, 2000), and lower termites (LT), which feed on wood and host cellulolytic flagellates in their gut (Inoue *et al*., 2000). Overall, the abundance of prokaryotic CAZymes was the highest in WF and the lowest in SF, while LT and FC fell between these two extremes (Table S7). This is consistent with the scarcity of lignocellulose in the diet of SF, which predominantly feed on the nitrogen-rich fraction of the soil, including microbial biomass and organic residues associated with clay particles (Ji and Brune, 2001, 2005; Ngugi *et al*., 2011; Ngugi and Brune, 2012). The intermediate abundance of prokaryotic CAZymes in FC and LT reflects their dependence on *Termitomyces* fungi for lignocellulose digestion (Poulsen *et al*., 2014) and on gut flagellates that encode for diverse cellulolytic enzymes (Yamin, 1981; Nishimura *et al*., 2020), respectively.

Task partitioning between gut prokaryotes and other symbionts –in which both partners participate in different steps of wood digestion and provide different sets of CAZymes– could be revealed from the gut metagenomes of LT and FC. Principal component analysis revealed that the prokaryotic CAZyme repertoire differs considerably among SF, LT, FC, and WF (Figure 3A). To characterize more accurately the contribution of termite gut prokaryotes to wood digestion, whenever possible, we identified the substrate of each 211 CAZymes (including 146 families and 65 subfamilies) present in more than 10% of termite species. We individually compared the abundance of these 211 CAZymes using phylogenetic ANOVA. We found that 178 comparisons were significantly different, and 177 comparisons remained significant after FDR corrections (Figure 3A, Table S7). Notably, we found that the combined seven GHs exclusively identified as cellulases were significantly depleted in LT as compared to other termite groups and were significantly depleted in FC and SF as compared to WF (Figure 2, Table S7). A similar pattern was found for the combined 29 GHs exclusively identified as hemicellulases, which were significantly more abundant in WF than in other termite groups (Figure 3A, Table S7). Therefore, the gut metagenomes of LT and FC appear to be depleted in prokaryotic GHs targeting cellulose as compared to WF, possibly reflecting task partitioning between termite gut prokaryotes and eukaryotic symbionts such as cellulolytic flagellates in LT and *Termitomyces* in FC. Task partitioning between gut prokaryotes and *Termitomyces* in FC was previously suggested for *Macrotermes natalensis* (Poulsen *et al*., 2014), with gut symbionts primarily participating to the final digestion of oligosaccharides and *Termitomyces* performing the breakdown of complex carbohydrates. In support of this hypothesis, several GHs, such as GH8, GH26, GH45, GH5_2, and GH53, largely depleted from the gut metagenomes of LT were highly expressed by the gut cellulolytic flagellates of *C. formosanus* (Nishimura *et al*., 2020), and were abundant in the gut metagenomes of WF. However, several GHs encoded by gut prokaryotes are also highly expressed by the gut cellulolytic flagellates of *C. formosanus* (*e.g.* GH13_8, GH36, GH3, GH92, GH133) (Nishimura *et al*., 2020). The extant of the complementarity between the CAZyme repertoires of gut flagellates and prokaryotes is therefore unclear and requires further investigation.

**Figure 3.**
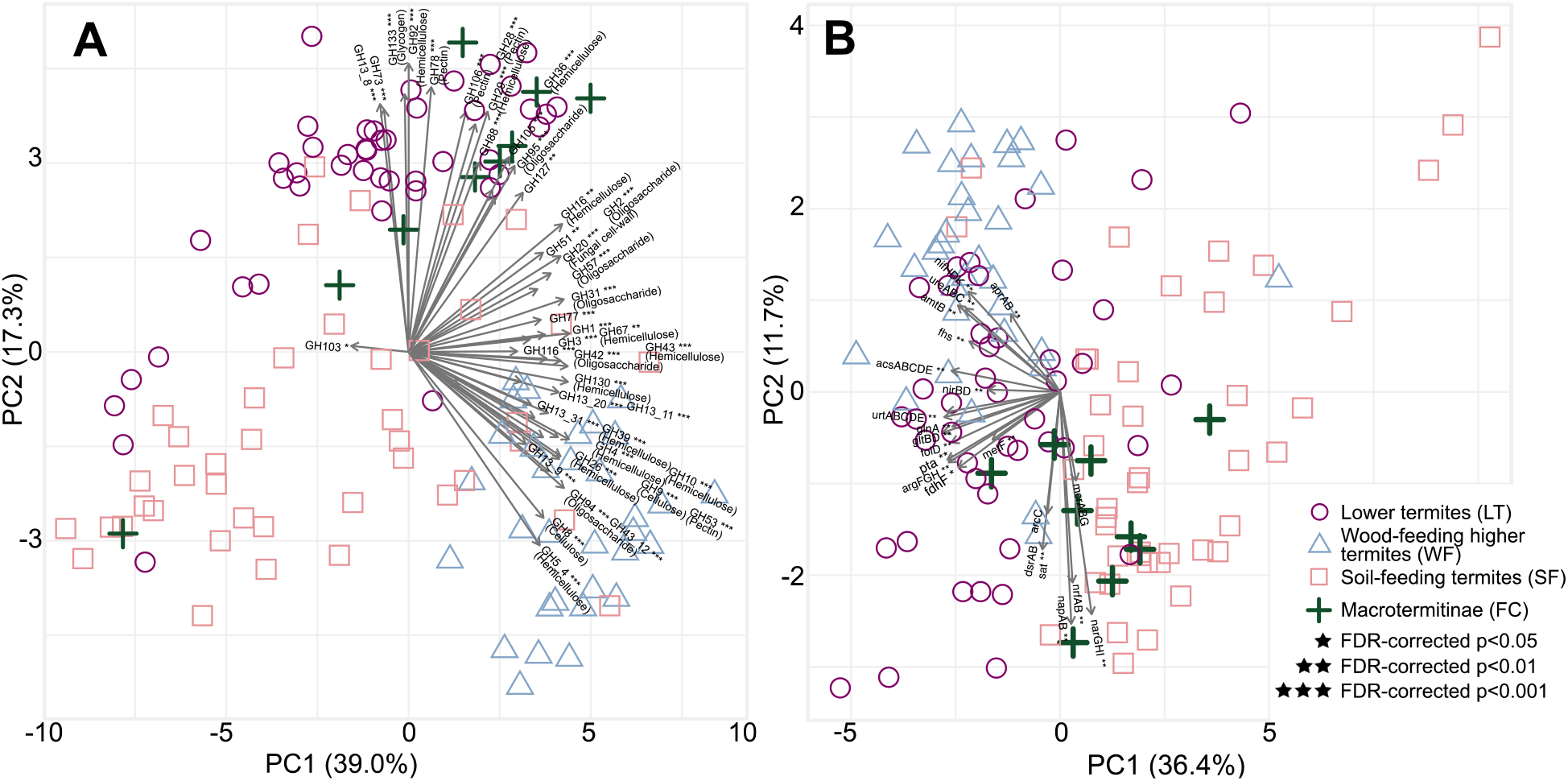
Principle component analysis (PCA) bi-plots showing the distribution of prokaryotic genes involved in lignocellulose digestion in the gut of termites. (A) PCA performed on the relative abundance of 346 CAZymes found in 129 gut metagenome assemblies. The 50 glycoside hydrolases (GHs) that contributed the most to separation of termite diets are plotted (see Table S7). (B) PCA inferred from relative abundance of metabolic genes involved in lignocellulose digestion after carbohydrate degradation. The symbols indicate host feeding habits. The species identity of each data point is available in Table S1. Asterisks indicate significant differences among the four termite groups at 5% false discovery rate (FDR, *p < 0.05; **p < 0.01; ***p < 0.001)

We next investigated the taxonomic origin of the prokaryotic CAZymes found in the same 129 whole gut metagenomes. We focused on the 19 GHs found in more than 10% of termite species and embedded in contigs longer than 5000 bps, allowing taxonomic annotation based on several genes. Contigs including genes with discordant taxonomic annotations potentially indicate horizontal gene transfers, as is common among bacteria (Ochman *et al*., 2000), and were removed. We found that Bacteroidota were a significant source of GH2, GH9, GH10, GH20, GH28, GH29, GH30, GH31, and GH130 in FC and LT, while, as previously described (Marynowska *et al*., 2020), they rarely encoded these GHs in non-Macrotermitinae Termitidae (WF and SF) (Figure 4, Table S8). In contrast, Fibrobacteres, which were very rare in LT, were a significant source of GH2, GH3, GH8, GH9, GH10, GH11, GH18, GH26, GH30, GH43, GH94, and GH130 in WF. Two other bacterial phyla, Spirochaetota and Firmicutes A, encoded most of the investigated GHs and were important contributors of GHs in WF (Figure 4, Table S8). Therefore, the primary contributors of GHs are distinct between lower and higher termites. These results are consistent with previous reports indicating a possible involvement of the ectosymbiotic Bacteroidota of some oxymonadid flagellates in cellulose and hemicellulose hydrolysis (Yuki *et al*., 2015; Treitli *et al*., 2019) in lower termites, while Fibrobacteres, Spirochaeota, and/or Firmicutes are major agents in cellulose and hemicellulose degradation in higher termites (Warneke *et al*., 2007; He *et al*., 2013; Tokuda *et al.,* 2018; Calusinska *et al*., 2020; Marynowska *et al.,* 2020). Our comprehensive analyses strongly indicate that the loss of cellulolytic flagellates in the ancestor of higher termites was accompanied by a major reworking of the cellulolytic bacterial communities, from Bacteroidota in LT to Fibrobacterota and Spirochaeota in WF and to Firmicutes in SF.

**Figure 4.**
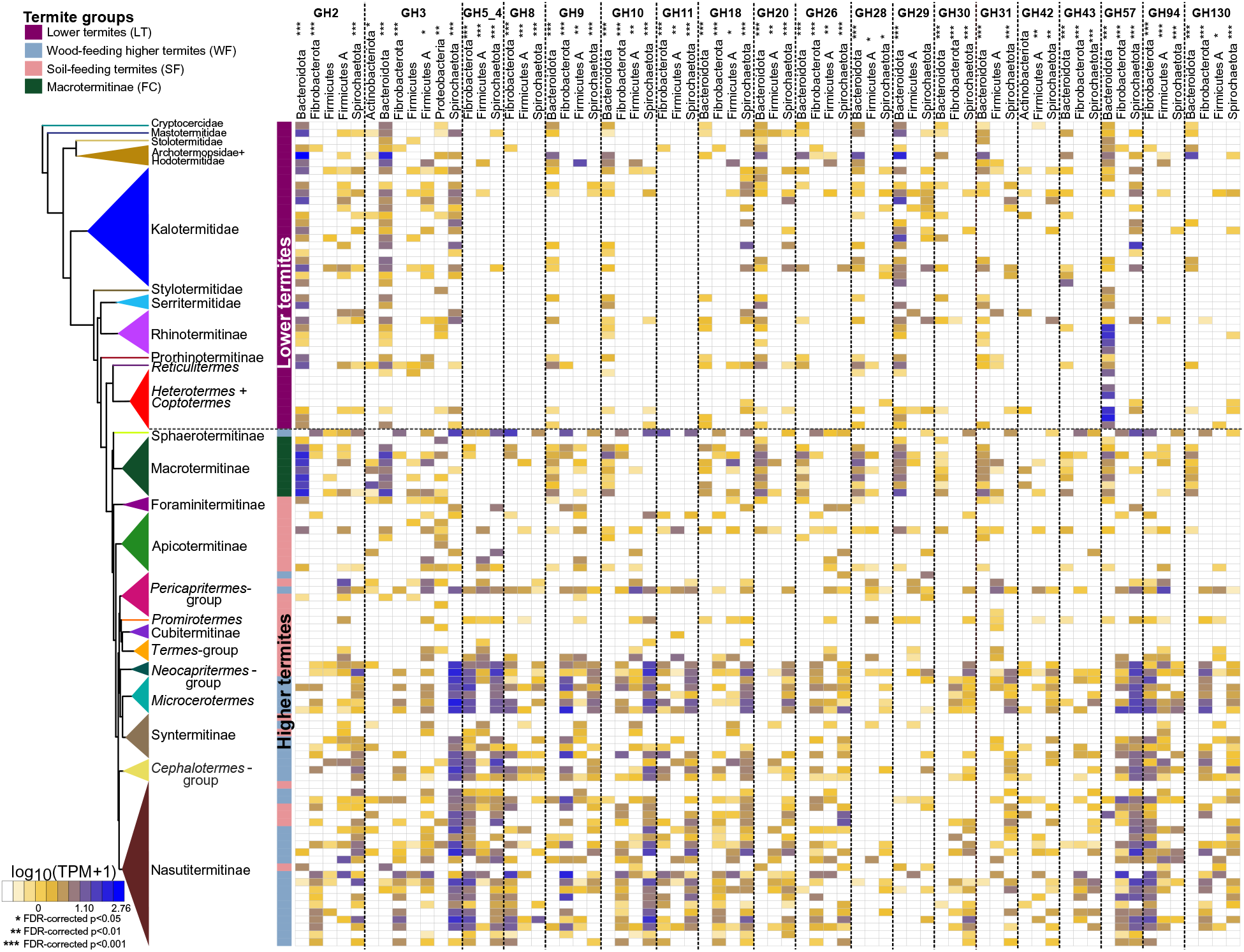
CAZyme families, and their taxonomic origin, for enzymes derived from contigs longer than 5000 bps and present in 10% of gut metagenomes. The color scale represents the log-transformed transcripts per million (TPM). The tree represents a simplified time-calibrated phylogenetic tree reconstructed using host termite mitochondrial genomes. Asterisks indicate significant differences among the four termite groups at 5% false discovery rate (FDR, *p < 0.05; **p < 0.01; ***p < 0.001).

CAZymes are often organized as polysaccharide utilization loci (PULs) that target complex polysaccharides (Terrapon *et al*., 2015). To search for PULs in our metagenomes, we reconstructed metagenome-assembled genomes (MAGs) by grouping contigs with similarities in sequence composition and depth of coverage. In total, we obtained 654 prokaryotic MAGs that ranged in completeness from 30% to 100% with <10% contamination for lineage-specific marker genes. We kept low quality MAGs, with completeness between 30% to 50%, as several such MAGs possessed complete pathways of interest (Figure S3, Table S9). The 654 MAGs included members of 16 phyla of bacteria and four phyla of archaea and included representatives of all major prokaryote phyla known to be present in termite gut. We found 128 PULs distributed across 130 MAGs, including 31 MAGs of Bacteroidota, 71 MAGs of Firmicutes, 13 MAGs of Proteobacteria, 12 MAGs of Spirochaetota, two MAGs of Actinobacteria, and one MAG of Verrucomicrobiota (Table S10). Sixteen PULs, found in 10 MAGs, had all the PUL components and mainly targeted lignocellulose components such as cellulose and xylan, and saccharides such as melibiose, alignate, and lactose. 107 PULs found in 74 MAGs encoded for more than one substrate but did not have all the PUL components, possibly reflecting the incompleteness of our MAGs or missing components nonessential for their activity, as experimentally demonstrated in the xylan utilization system (*Xus*) of a Bacteroidota associated with *Pseudacanthotermes* (Wu, 2018). Altogether, our data provide an overview of the PUL distribution in termite gut microbes.

### Reductive acetogenesis in termite gut

The fermentation of wood fibers by the termite gut microbiota produces mostly acetate, which is used by the termite host, but also H_2_ and CO_2_ (Hungate, 1939; Brune, 2014). Most of the H_2_ is used to produce additional acetate by the reduction of CO_2_ (Breznak and Switzer, 1986; Brauman *et al*., 1992; Pester and Brune, 2007). We focused on the genes of seven enzymes of the Wood-Ljungdahl pathway (WLP) of reductive acetogenesis that are present in all acetogens from termite guts identified to date, namely formate dehydrogenase H (*fdhF*), formate-tetrahydrofolate ligase *(fhs),* methylenetetrahydrofolate dehydrogenase *(folD),* 5,10-methylenetetrahydrofolate reductase *(metF)*, acetyl-CoA synthase (*acsABCDE)*, phosphotransacetylase (*pta*), and acetate kinase (*ack*), which are essential to operate the bacterial WLP (Schuchmann and Müller, 2014). We compared the relative abundance of these markers across the 129 whole gut metagenomes used for previous analyses and found a significant phylogenetic autocorrelation signal with the termite phylogenetic tree for five of the seven enzymes, two of which remain significant after FDR correction (*fdhF* and *acsABCDE*) (Figure 5, Table S11). Together with the five other enzymes, which also occur in many other bacteria, the simultaneous presence of *fdhF* and *acsABCDE* is a strong predictor for the distribution of reductive acetogenesis across the termite phylogenetic tree.

**Figure 5.**
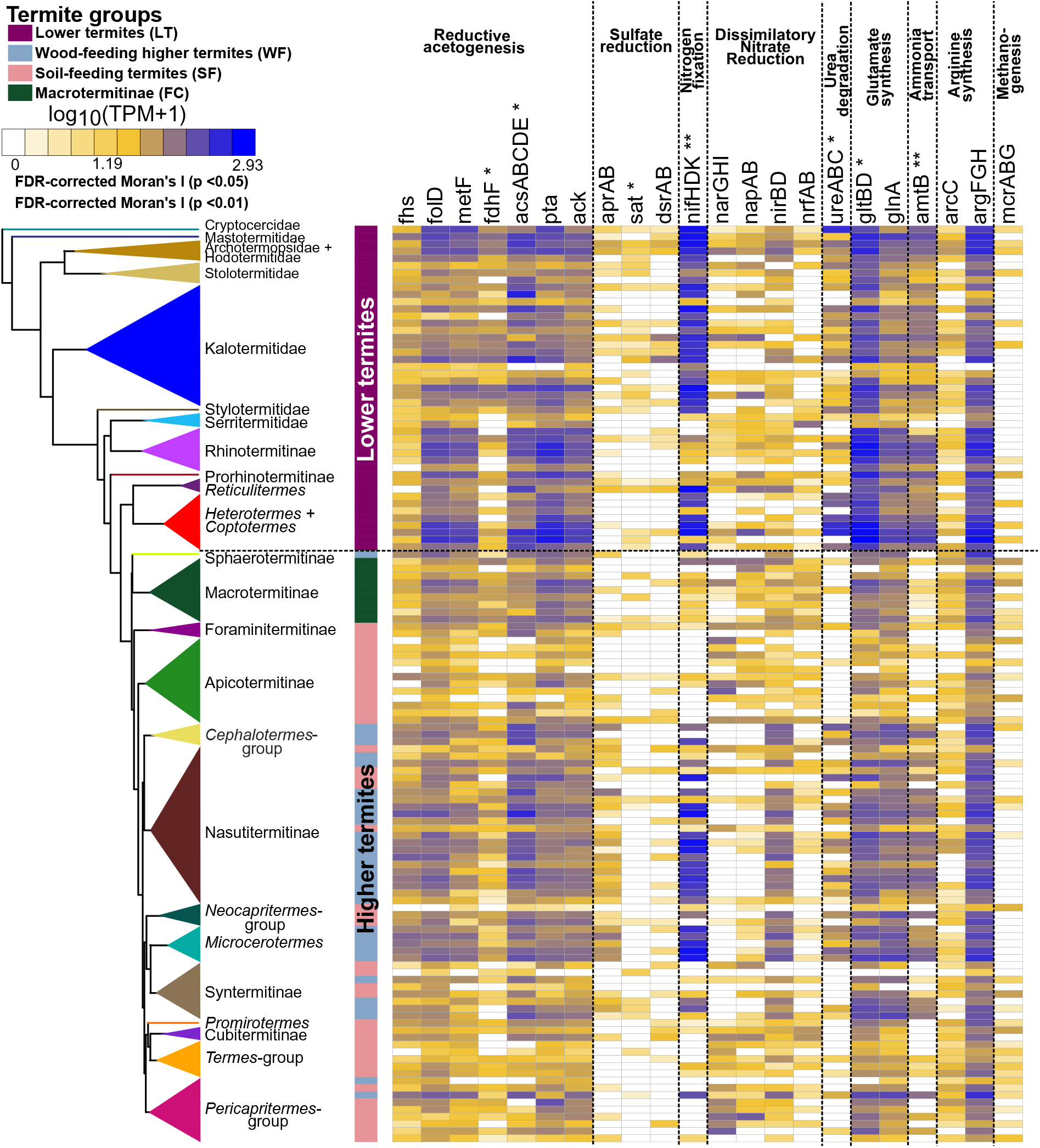
Relative abundance of prokaryotic genes belonging to metabolic pathways involved in the final steps of the lignocellulose digestion in the gut of termites. The color scale represents the logarithm of transcripts per million (TPM). The tree represents a simplified time-calibrated phylogenetic tree reconstructed using host termite mitochondrial genomes. Full names of the gene families and their corresponding KEGG IDs are available in Table S11.

The seven enzymes encoded by all acetogens significantly differed in relative abundance among the four termite groups. They were generally more abundant in LT and WF than in FC and SF (Figure 3B, Table S11). These analyses are in agreement with previous studies that measured the potential rates of acetogenesis in a smaller set of termite species, and corroborate the hypothesis that reductive acetogenesis is mostly associated with a diet of wood and is less important in fungus-cultivating Macrotermitinae and in soil-feeding lineages (Brauman *et al*., 1992; Tholen and Brune, 1999).

To determine the identity of the acetogens, we searched each MAG for the genes of the seven enzymes associated with reductive acetogenesis. We found 44 MAGs associated with six termite families and *Cryptocercus* that encoded at least five of the seven enzymes, but none of these MAGs contained the complete set of genes (Table S12, Figure 6A). In addition to formate dehydrogenase H (*fdhF*), we also searched for the genes encoding [FeFe] hydrogenase Group A4 (*HydA*) and the iron-sulfur cluster proteins (*HycB3*, *HycB4*), the other subunits of the hydrogen-dependent CO_2_ reductase (HDCR) complex catalyzing the first step of CO_2_ reduction to formate (Schuchmann and Müller, 2012; Ikeda-Ohtsubo *et al*., 2016). Two MAGs lacked *fdhF* but contained all other genes of the WLP and the HDCR complex (Table S12, Figure 6A). These MAGs belonged to the Desulfobacterota family *Adiutricaceae*, which comprises the putatively acetogenic *Candidatus* Adiutrix intracellularis, a flagellate endosymbiont from the archotermopsid *Zootermopsis*, and numerous uncharacterized representatives from other lower and higher termites (Ikeda-Ohtsubo *et al*., 2016). Like *Ca.* Adiutrix intracellularis, none of the four MAGs encoded a sulfate reduction pathway. They were found in the rhinotermitid *Dolichorhinotermes* and in the higher termite *Microcerotermes*, indicating that the putatively free-living members of *Adiutricaceae* from higher termites (which lack gut flagellates) are also acetogenic.

**Figure 6.**
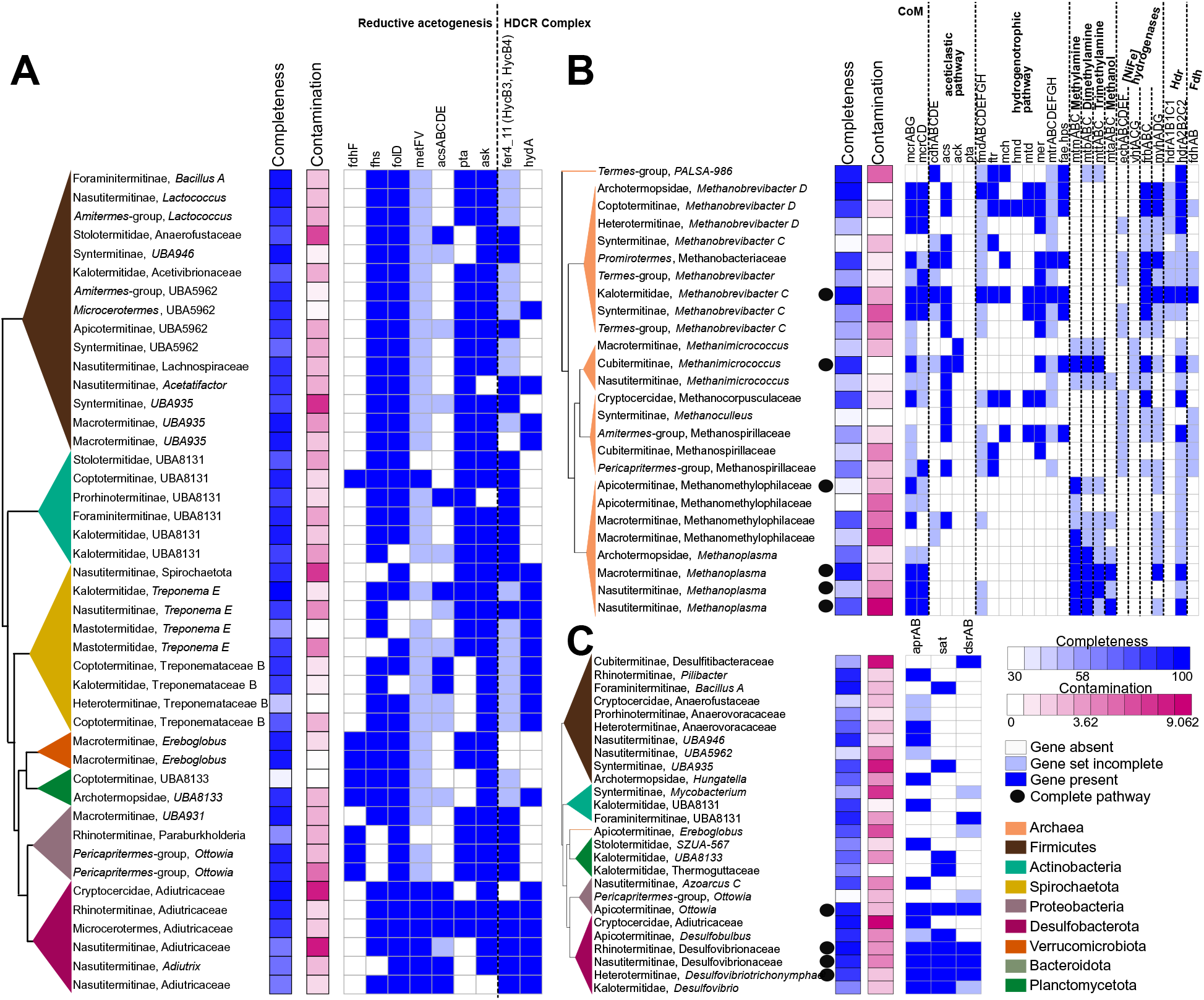
Metabolic pathways involved in the final steps of lignocellulose digestion found in gut metagenome assembled genomes (MAGs) reconstructed in this study. (A) Genes involved in reductive acetogenesis, (B) methanogenesis, and (C) sulfate reduction found in MAGs. The trees represent simplified maximum likelihood phylogenetic trees of the MAGs reconstructed using 43 single-copy marker genes. MAG completeness and contamination, based on CheckM analyses, is shown beside the tree. Dark blue squares indicate gene presence, light blue squares indicate that incomplete gene sets, and open squares indicate gene absence. Detailed information on the gene families and their KEGG IDs are available in Tables S12, S14, and S15.

Because none of the other MAGs encoded a complete WLP, we could not unambiguously attribute acetogenic status to any other prokaryote lineage. Considering the high rates of reductive acetogenesis in many lower and higher termites, particularly the wood-feeding species (Brauman *et al*., 1992), this may be explained either by the incompleteness of our MAGs or the failure to assemble any genomes of the populations responsible for the acetogenic activity. Based on the low free energy yields of both reductive acetogenesis and methanogenesis, it has been speculated that the proportion of (hydrogenotrophic) acetogens among the prokaryotic community in termite hindguts may be as low as that of (hydrogenotrophic) methanogens (Loh *et al*., 2021). The problem of genome assembly from low abundance populations would be exacerbated by a high species diversity among members of a particular metabolic guild. Alternatively, the absence of a complete reductive acetogenesis pathway among our MAGs may be genuine. This could be the case among the MAGs assigned to the family Treponemataceae B. Although the first isolate of this lineage is a homoacetogen with a complete WLP (Leadbetter et al., 1999), none of the other species isolated to date are acetogenic (Song *et al*., 2021). With the exception of *Treponema primitia* (Graber *et al*., 2004), *Candidatus* Treponema intracellularis (Ohkuma *et al*., 2015), and *Candidatus* Adiutrix (Ikeda-Ohtsubo *et al*., 2016), the identity of the populations responsible for reductive acetogenesis in termite guts, including the putatively acetogenic *Candidatus* Termitimicrobium (Bathyarcheia; Loh et al., 2021) remains open to speculation.

### Methanogenesis in termite gut

The methanogenic archaea present in the gut of termites consume a large fraction of H_2_ and are responsible for 3% of global methane emissions (Brune, 2018, 2019). We searched the 129 gut metagenomes used in earlier analyses for genes that are part of methanogenesis pathways. Because of the low abundance of Archaea in termite guts (Brune 2019; Loh *et al*., 2021), the abundance of genes involved in methanogenesis was often near, or below, our detection threshold. As a consequence, we were unable to analyze each gene independently, but instead calculated the Moran’s I index using the abundance of genes encoding the methyl-coenzyme M reductase complex (*mcrABG*), which catalyzes the final step of methanogenesis (Evans *et al*., 2019), and found no autocorrelation signal with the termite phylogenetic tree (Figure 5, Table S11).

We compared the abundance of *mcrABG* among the four termite groups and found no significant differences (Figure 3B, Table S11). However, this lack of significance probably reflects the low abundance of archaeal reads in our assemblies, rather than an actual uniformity of methanogenesis pathways across termites, as methane emission rates are known to be diet-related and particularly high in species feeding on soil (*e.g.*, Brauman *et al*., 1992; Bignell and Eggleton, 1995; Bignell *et al*., 1997; Sugimoto *et al*., 1998).

We searched our gut metagenomes for operons encoding *mcrABG*, and found 14 operons, belonging to four methanogenic archaeal orders, *Methanomassiliicoccales*, *Methanobacteriales*, *Methanomicrobiales*, and *Methanosarcinales*, derived from the gut metagenomes of 14 termite species, including four of the eight families of LT, and five of the nine subfamilies of Termitidae (Table S13). All *mcrABG* operons of LT were classified to *Methanobacteriales*, which is in agreement with previous reports on the prevalence of *Methanobacteriales* in LT (Brune, 2019). An exception was found in the gut metagenome of *Porotermes quadricollis*, which yielded an *mcrABG* operon from *Methanomethylophilaceae* (order *Methanomassiliicoccales*). This is unusual, because members of this order are frequently encountered in higher termites and millipedes (Paul *et al*., 2012) but have been detected only once in the lower termite *Reticulitermes speratus* (Shinzato *et al*., 2001).

Next, we analyzed the methanogenic capacities of 26 MAGs of Archaea reconstructed from the gut metagenomes of 23 termite species from four termite families and the cockroach *Cryptocercus*. Only 13 MAGs belonging to *Methanomicrobiales*, *Methanobacteriales*, *Methanosarcinales*, and *Methanomassiliicoccales* encoded the *mcrABG* complex, indicating that the assemblies are incomplete (Figure 6B, Table S14). Five of these 13 MAGs possessed complete pathways for methylotrophic methanogenesis and one MAG possessed complete pathways for hydrogenotrophic methanogenesis (Figure 6B). The five MAGs showing genomic evidence of methylotrophic methanogenesis included one MAG of *Methanosarcinales* (genus *Methanimicrococcus*) and four MAGs of *Methanomassiliicoccales*, including three MAGs classified to genus *Methanoplasma* and one MAG classified to family *Methanomethylophilaceae*. Only two MAGs of *Methanoplasma* encoded a methanol:coenzyme M methyltransferase (*mtaABC*) complex, which is required for growth on methanol and typical for all members of this lineage (Lang et al., 2015), and only one of the MAG of *Methanosarcinales* and one MAG of *Methanoplasma* encoded a complete heterodisulfide reductase complex (*HdrA2B2C2/mvhADG*) present in most methanogens (Thauer *et al*., 2008; Buckel and Thauer, 2013), underscoring the incompleteness of the MAGs. The same was true for hydrogenotrophic methanogenesis, for which only one MAG belonging to *Methanobacteriaceae* (genus *Methanobrevibacter C*) possessed most of the genes required for the reduction of CO_2_ to methane, including a heterodisulfide reductase (*HdrABC/mvhADG*) complex, an iron-sulfur flavoprotein along with a F420-independent hydrogenase (*Fdh*), and a F420 reducing hydrogenase (*FrhABC*) (Figure 6B, Table S14). The absence of aceticlastic methanogens is in agreement with previous reports (Brune, 2018, 2019). Overall, our results highlight the diversity of methanogens found in termite guts, and the diversity of the pathways they use.

### Sulfate-reducing prokaryotes

Sulfate-reducing bacteria are potential H_2_-consumers in the gut of termites (Brauman *et al*., 1990; Kuhnigk *et al*., 1996; Dröge *et al*., 2005) (Figure 5). However, sulfate concentration is low in termite gut, as is H_2_ consumption by sulfate-reducing bacteria (Dröge *et al*., 2005; Brune and Ohkuma, 2011). We found all the genes of the dissimilatory sulfate reduction pathway, namely, the two subunits of adenylylsulfate reductase (*aprA* and *aprB*), sulfate adenylyltransferase (*sat*), and dissimilatory sulfite reductase (*dsrAB*), in six out of eight lower termite families, all the higher termite subfamilies, and *Cryptocercus*. The abundance of *aprAB* and *sat* were significantly correlated with the termite phylogenetic tree, and the correlation remained significant after FDR correction for *sat* (Figure 5, Table S11).

Comparisons of the four termite groups showed that the abundance of *aprAB* was significantly higher in WF than in SF and the abundance of *sat* was significantly higher in LT than WF and SF (Figure 3B, Table S11). While sulfate reducers have been isolated from the guts of LT, FC and SF (Brauman *et al*., 1990; Kuhnigk *et al*., 1996), we found metagenomic evidence that sulfate reduction is also prevalent in WF.

Next, we analyzed the sulfate-reducing capabilities of our 654 MAGs and found a complete pathway for dissimilatory sulfate reduction in four MAGs (Figure 6C, Table S15). Three of these MAGs, found in the termites *Parrhinotermes*, *Reticulitermes*, and *Tumulitermes*, were assigned to *Desulfovibrionaceae* (Desulfobacterota), which are common in the termite gut and generate energy via sulfate respiration (Sato *et al*., 2009; Kuwahara *et al*., 2017). Of note, the fourth MAG, retrieved from the gut metagenome of the apicotermitine *Heimitermes laticeps*, belonged to the Proteobacteria family *Burkholderiaceae*, a bacterial family that was, prior to this study, largely unreported from termite guts, and that is abundant in Apicotermitinae and in the termite clade that includes the Cubitermitinae, the *Pericapritermes*-group, and the *Termes*-group. The evidence for dissimilatory sulfate reduction in *Burkholderiaceae* termite guts suggest that the capacity for sulfate respiration is more widely distributed than expected.

### Nitrogen recycling by termite gut prokaryotes

Because the content of nitrogen in wood is low, termites have evolved mechanisms of nitrogen conservation. The termite gut microbiota contributes to the nitrogen metabolism of its host by recycling nitrogen (Breznak, 2000; Hongoh, 2011). Like most insects, termites convert waste products from nitrogen metabolism into uric acid, but, unlike other insects, the gut prokaryotes of termites degrade uric acid into ammonia, which is subsequently assimilated by the gut microbiota (Brune, 2014). We searched the 129 metagenomes used for previous analyses and found only few genes possibly involved in uric acid degradation, including 11 *aegA* (a putative oxidoreductase suspected to be involved in uric acid degradation by Enterobacteriaceae (Iwadate and Kato, 2019)) in six termite species. Since the uricolytic prokaryotes isolated from termite guts are strict anaerobes (Potrikus and Breznak, 1980, 1981; Thong-On *et al*., 2012), it is likely that they use alternative, so far unknown, pathways. Termite tissues reportedly lack uricase activity (Potrikus and Breznak, 1981), but when we examined the transcriptomes of 53 termite species generated by Buček *et al*. (2019), we found evidence for the expression of a gene encoding urate oxidase in 20 termite species belonging to four termite families (Figure S4). This indicates that termites should be able to carry out the first step of uric acid degradation. However, the extent of the contribution of the termite host to uricolysis and the identity of the uricolytic prokaryotes and their catabolic pathways remain unknown.

The metagenomes of all termite families included numerous prokaryotic genes from other pathways involved in the production of ammonia (Figure 5, Table S11), including ureases (*ureABC*), which degrade urea into ammonia (Hongoh and Ohkuma, 2010; Ohkuma *et al*., 2015), and some of the genes of the dissimilatory nitrate reduction pathway (*narGHI*, *napAB*, *nrfAB*), which convert nitrate into ammonia. Among those, the abundance of *ureABC* genes significantly correlated with the termite phylogenetic tree after FDR correction (Figure 5, Table S11). We also found in the metagenomes of all termite families genes from pathways involved in amino acid biosynthesis from ammonia, including glutamine synthetase (*glnA*) and glutamate synthase (*gltBD*), the genes involved in the synthesis of glutamate from ammonia, and carbamate kinase (*arcC*), ornithine carbamoyltransferase (*argF*), argininosuccinate synthase (*argG*) and argininosuccinate lyase (*argH*), the genes involved in arginine biosynthesis from ammonia (Yan 2007). The abundance of *gltBD* correlated with the termite phylogenetic tree after FDR correction (Figure 5, Table S11). Therefore, the termite phylogeny is a good predictor of the enteric abundance of some of the prokaryotic genes involved in ammonia metabolism in termites.

We compared the four termite groups using the relative abundance of the nitrogen-recycling genes and found that the abundance of *ureABC* differed among termite groups, with the gut metagenomes of LT and WF significantly enriched in *ureABC* as compared to those of SF and FC (Figure 3B, Table S11). In contrast, the abundance of some of the genes of the dissimilatory nitrate reduction pathway, such as *napAB* and *narGHI*, was significantly reduced in the gut metagenomes of WF compared to SF and FC (Figure 3B, Table S11). This suggests that the high rates of nitrate ammonification previously found in two soil-feeding species (Ngugi and Brune, 2012) is a characteristic that all soil-feeding termites share with fungus-cultivating termites. We also found that *gltBD* was significantly enriched in LT as compared to other termite groups, while *argFGH* was significantly enriched in LT and WF as compared to SF (Figure 3B, Table S11). The low abundance of genes involved in ammonia assimilation in soil-feeding termites is likely linked to their diet, which includes soil peptidic residues (Ji and Brune, 2001, 2005).

Next, we searched our 654 MAGs to determine the taxonomic identity of the prokaryotes involved in nitrogen recycling. Six MAGs possessed the three ureases *ureABC*, thence encoded enzymes to convert urea into ammonia, and 15 MAGs included a complete dissimilatory nitrate reduction pathway that convert nitrate into ammonia. All these MAGs belonged to diverse lineages of Proteobacteria and Campylobacterota (order *Campylobacterales*), except for one MAG of Firmicutes (genus *Bacillus*) found in *Foraminitermes rhinoceros* and endowed with *ureABC*, *narGHI* and *nirBD* (Figure 7A, Table S16). We also found numerous MAGs capable of ammonia assimilation into glutamate and arginine, indicating that ammonia is an important source of nitrogen for many termite gut prokaryotes. 91 MAGs possessed *glnA* and *gltBD* for glutamate biosynthesis from ammonia, while 26 MAGs possessed the four genes *arcC*, *argF*, *argG*, and *argH* for arginine biosynthesis from ammonia via the urea cycle, including 12 MAGs that also contained the glutamate biosynthesis pathway (Figure 7A, Table S16). 66 MAGs encoding glutamate biosynthesis genes, and 15 MAGs with arginine biosynthesis genes, also possessed the ammonium transporter *Amt*. These MAGs belonged to ten phyla, including 19 MAGs of Proteobacteria from six families,18 MAGs of Bacteroidota, of which eight belonged to the family *Azobacteroidaceae*, ten MAGs of Actinobacteria of three families, eight MAGs of the Spirochaetes family *Treponemataceae* B, eight MAGs of Firmicutes, six MAGs of *Campylobacterota*, five MAGs of Firmicutes A, three MAGs of Planctomycetota, three MAGs of Desulfobacterota, and one MAG of Verrucomicrobia (Figure 7A, Table S16). Therefore, a great many bacterial lineages contribute to the nitrogen metabolism of their termite hosts.

**Figure 7.**
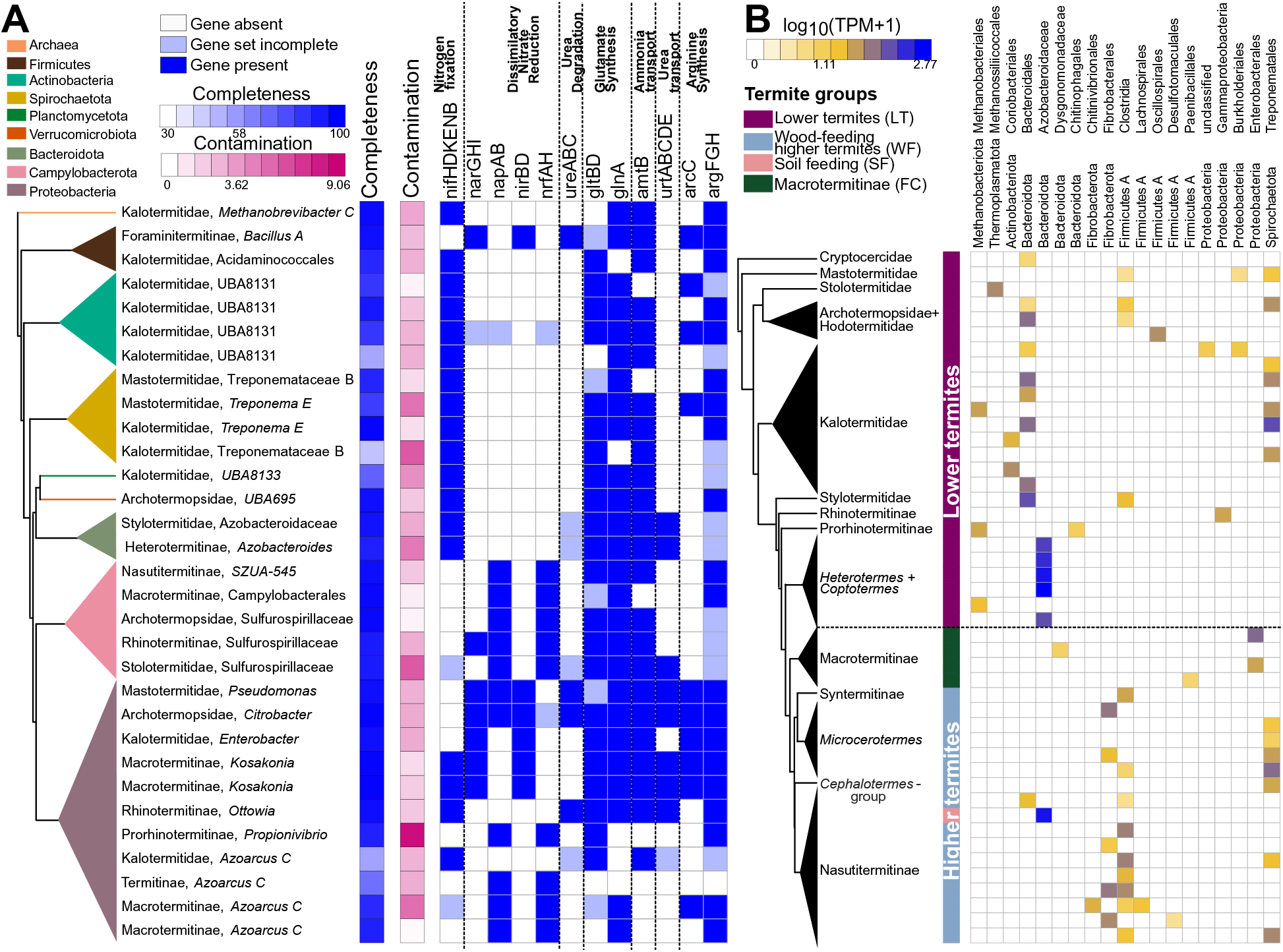
Nitrogen metabolism in the gut of termites. (A) Metagenome-assembled genomes (MAGs) with complete nitrogen fixation or dissimilatory nitrate reduction pathways. All pathways potentially involved in the nitrogen metabolism, namely nitrogen fixation, dissimilatory nitrate reduction, ureases, glutamate metabolism, ammonia transport, urea transport, and arginine metabolism are represented. The tree represents a simplified maximum likelihood phylogenetic tree of the MAGs inferred from 43 marker genes. Completeness and contamination of MAGs, based on CheckM analysis, are shown beside the tree. Dark blue squares indicate gene presence, light blue squares indicate that incomplete gene sets, and open squares indicate gene absence. (B) Abundance of NifHDK operons (*nifHDK*, *vnfHDK*, or *anfHDK*) present in contigs longer than 5000bps across gut metagenomes. The color scale represents the log-transformed transcripts per million (TPM). The tree represents a simplified time-calibrated phylogenetic tree reconstructed using host termite mitochondrial genomes.

### Nitrogen fixation by termite gut prokaryotes

Many species of wood-feeding termites host dinitrogen-fixing prokaryotes in their gut, which compensate for the low nitrogen content of wood (Breznak, 2000). They fix nitrogen with either the molybdenum-dependent (Nif), vanadium-dependent (Vnf), or iron-only alternative nitrogenases (Anf) (Ohkuma *et al*., 1999; Yamada *et al*., 2007; Inoue et al. 2015). We found gene homologs for the structural subunits of these nitrogenases (collectively referred to as *nifDHK*) in all metagenomes of termite families and in *Cryptocercus* (Figure 5). Their abundance significantly correlated with the termite phylogeny after FDR correction (Figure 5, Table S11), as was the case for several other pathways involved in nitrogen economy. There were significant differences among termite groups, with the nitrogenase reads in the gut metagenomes of non-FC wood-feeders (LT and WF) being 24.4-fold more abundant than in SF and 20.2-fold more abundant than in FC (Figures 3B, 5, Table S11). This is in line with the higher rate of N_2_ fixation measured in LT and WF than in SF and FC (Yamada *et al*., 2007), and reflects the high amount of nitrogen present in soil and fungi, making the energy-demanding process of N_2_ fixation unnecessary (Brune and Ohkuma, 2011; Hongoh, 2011).

To identify the diazotrophs present in the gut of termites, we taxonomically classified contigs longer than 5000 bps that contained the six genes present in all diazotrophs, *nifDHK* (which encode nitrogenase), and *nifB*, *nifE*, *nifN*, which encode proteins involved in nitrogenase biosynthesis (Dos Santos *et al*., 2012). We identified 15 contigs matching these criteria in the gut metagenomes of 12 termite species, representing five of the nine termite families (Table S17). These contigs were assigned to diverse prokaryote lineages, including nine contigs of diverse Bacteroidota, three contigs of the Spirochaetota order *Treponematales*, two contigs of Proteobacteria family *Enterobacteriaceae*, and one contig of the archaeal genus *Methanobrevibacter*. We carried out the same analyses on our MAGs and found 18 MAGs that contained a *nifHDKBEN* cluster, including seven MAGs that belonged to phyla not represented in the contigs >5000 bps. Among these seven MAGs, there were four MAGs of the Actinobacteriota family *UBA8131*, one MAG of the Planctomycetota family *Thermoguttaceae*, one MAG of the Verrucomicrobiota family *Chthoniobacteraceae*, and one MAG of the Firmicutes C order *Acidaminococcales* (Figure 7A, Table S16). Therefore, the taxonomy of diazotrophs found in our termite species set corroborates previous evidence that termites host diverse communities of diazotrophs in their guts (Ohkuma *et al*., 2001; Yamada *et al*., 2007; Desai and Brune, 2012).

We next investigated the taxonomic distribution of diazotrophs across termites. We focused on contigs longer than 5000 bps that included genes with concordant taxonomic annotation and that contained a *nifHDK* operon (Figure 7B, Table S18). In lower termites, the dominant diazotroph was an undescribed Bacteroidota allied to an ectosymbiont of the *Cryptocercus* gut flagellate *Barbulanympha* (Tai *et al*., 2016). This undescribed Bacteroidota was found in three of the eight families of LT. It was also largely absent from the gut metagenomes of *Coptotermes* and *Heterotermes*, which harbor the flagellate endosymbiont *Candidatus* Azobacteroides as the main diazotroph (Hongoh *et al*., 2008). The diazotrophs of Termitidae belonged to various phyla. Notably, we found the N_2_-fixing *Candidatus* Azobacteroides in the nasutitermitine *Coatitermes* (which lacks gut flagellates), and a N_2_-fixing *Treponematales* in *Mastotermes*, highlighting that the dominant lineages of diazotrophs in particular termite lineages are also harbored at a low abundance by unrelated species of termites (Figure 7B, Table S18). Therefore, our results indicate that the phylogenetic position of termite species determined, at least partly, the taxonomy of their dominant diazotrophs.

## CONCLUSION

The metagenomics and metatranscriptomics surveys of termite guts carried out so far targeted a limited number of termite species (*e.g.* Warnecke *et al*., 2007; He *et al*., 2013; Liu *et al*., 2018; Tokuda *et al*., 2018; Marynowska *et al*., 2020), and thus did not permit an investigation of how the gut microbiome of these social roaches has been evolving in term of function and composition since termite origin, ∼150 Million years ago. To address this issue, and to provide a global picture of the taxonomic and functional composition of the termite gut microbiome, we generated gut metagenomes for a comprehensive set of 145 termite species. The analyses of this dataset revealed that: (1) gut prokaryotic genes involved in the main nutritional functions are generally present across termites, suggesting these genes were already harbored by the common ancestor of modern termites; (2) the termite phylogenetic tree is largely predictive of the gut bacterial community composition and the nutritional function they exert; and (3) the acquisition of a diet of soil was accompanied by a change in the stoichiometry of genes and metabolic pathways involved in important nutritional functions rather than by the acquisition of new genes and pathways.

The analyses of our 146 gut metagenomes indicated that prokaryotic CAZymes, genes of the reductive acetogenesis, sulfur reduction, and methanogenesis pathways, and genes involved in nitrogen fixation and recycling, are present across the nine termite families. Therefore, the nutritional functions previously known to be performed by the gut prokaryotic symbionts of particular termite species (*e.g.* Warnecke *et al*., 2007; Calusinska *et al*., 2020) are performed in the gut of all termites. These results strongly suggest that the gut prokaryotes performing important nutritional functions were already harbored by the common ancestor of modern termites. Following this scenario, the ancestor of termites did not only acquire their charismatic gut cellulolytic flagellates (Nalepa, 1991), but also acquired several bacterial and archaeal lineages that make up a sizable fraction of the gut microbiota of modern termite species. In support of this hypothesis, many termite gut bacteria phylotypes form monophyletic groups present in the gut of various termite families and distantly related to bacteria found in other environments, such as in the guts of other animals, including cockroaches (Bourguignon *et al*., 2018). Therefore, as the cockroach-like ancestor of termites evolved wood-feeding, it is likely that it recruited facultative gut microbes able to degrade wood and participate in the nitrogen economy as essential gut symbionts.

Our analyses indicate that the phylogenetic position of termite species is partly predictive of the functions of gut bacterial communities. This is best illustrated by CAZymes whose abundance often correlated with the termite phylogenetic tree. Correlation with the termite phylogenetic tree, however, was not found for some genes, such as the *mcrABG* genes of the methanogenesis pathway, the genes of sulfate reduction pathway, and the genes of the dissimilatory nitrate reduction pathway. Whether this lack of correlation is genuine, or whether it reflects insufficient depth of sequencing, is unclear and requires further study. In any case, our results indicate that the correlation found between the phylogenetic tree of termites and their gut bacterial and protist communities (Rahman *et al*., 2015; Tai *et al*., 2015) are also found for some gut microbial functions.

The comparison of four termite groups, soil-feeding Termitidae (SF), fungus-cultivating Macrotermitinae (FC), non-Macrotermitinae wood-feeding Termitidae (WF), and lower termites (LT), reveals that genes and metabolic pathways important to termites are present in all termite species, but their abundances vary among groups. Notably, the gut metagenomes of SF possessed on average fewer CAZymes, nitrogenases, reductive acetogenesis, and sulfate-reducing genes than the gut metagenomes of other termite groups. Therefore, as pointed out by Marynowska *et al*. (2020), the gut prokaryote communities of SF retain important carbohydrate metabolism capabilities. Nevertheless, our dataset clearly indicate that these abilities are much reduced in soil-feeders compared to wood-feeders. Overall, our results support the idea that the acquisition of soil-feeding was accompanied by changes in the abundance of the gut prokaryote metabolic pathways important to termite nutrition.

## ACKNOWLEDGMENTS

We thank the DNA Sequencing Section and the Scientific Computation and Data Analysis Section of the Okinawa Institute of Science and Technology Graduate University, Okinawa, Japan, for assistance with sequencing and for providing access to the OIST computing cluster, respectively.

## AUTHORS’ CONTRIBUTIONS

J.A., Y.K., and T.B. conceptualized the experiments and approach. J.S., P.S. Y.R., Y.C.P., K.Y.K., D.S.D., G.T., and T.B. collected the samples. J.A and C.C performed the lab experiments. J.A., Y.K., and T.B. designed the data analyses. J.A. and Y.K. performed the bioinformatics analyses. A. Bucek examined the urate oxidase function in termite transcriptomes. J.A. and T.B. wrote the paper, with significant contribution from A. Brune, V.H., and G.T. All authors read and accepted the final version of this manuscript.

## CONSENT FOR PUBLICATION

Not applicable.

## DATA AVAILABILITY

Raw sequence data generated in this study have been deposited on MG-RAST (https://www.mg-rast.org/mgmain.html?mgpage=project&project=mgp100619) (see Table S1 for individual IDs). MAGs generated in this study are available on Figshare (https://doi.org/10.6084/m9.figshare.17031674.v1). The scripts used in this study are available on github (https://github.com/oist/EGU-The-functional-evolution-of-termite-gut-microbiota).

## ETHIC APPROVAL

Not applicable.

## METHODS

### Sample collection

We collected a total 145 termite samples and one sample of the cockroach *Cryptocercus kyebangensis* (Table S1, Figure S1). These samples were representative of the global termite diversity. All samples were preserved in RNA-later® and stored at −80°C until DNA extraction.

### DNA extraction and sequencing

Genomic DNA extraction was performed on the whole guts of five workers using the NucleoSpin Soil kit (Macherey-Nagel) according to manufacturer’s protocol. Library preparation was performed using the KAPA Hyperplus Kit, which is based on a unique dual tag indexing approach that minimizes the effects of index hopping. Libraries were either PE250-sequenced on the Illumina HiSeq2500 platform or PE150-sequenced on the Illumina HiSeq4000 platform (Table S1).

### Data filtering and assembly of metagenomic reads

Raw reads were filtered based on their quality. Reads with average Phred quality score below 30 were removed using Trimmomatic v 0.33 (Bolger *et al*., 2014). The “SLIDINGWINDOW” was set to “4:30” to trim low quality bases (Phred quality score below 30) from the 3’ end of the reads. We removed the 16 base pairs at the 5’ end of each read using the “HEADCROP” option because we observed over-represented k-mers in this region of the reads. Reads shorter than 50bps were removed.

The quality-controlled reads were assembled into contigs using SPAdes v 3.11.1 (Nurk *et al*., 2017) with the “meta” option and k-mer sizes of 21, 31, 41, 51, 71. The assembly quality was checked using the “metaquast” option of QUAST v 3.1 (Quality Assessment for Genome Assemblies) based on weighted median contig size (N50) (Gurevich *et al*., 2013) and percent of reads mapped to the contigs (Langmead *et al*., 2012; Papudeshi *et al*., 2017). Only the reads mapped to prokaryotic contigs were examined in this study (see the *taxonomic annotation* and *functional annotation* sections below).

### Termite phylogenetic tree reconstruction

We build a phylogenetic tree of termites using mitochondrial genomes retrieved from metagenome assemblies. Mitochondrial contigs derived from termites were identified using BLAST search (sequence length >5000 and percent identity >90) (Altschup *et al*., 1990) against previously published whole mitochondrial genomes of termites (Bourguignon *et al*., 2015, 2016, 2017; Wang *et al*., 2019). Mitochondrial genomes were complete, or near-complete, in most cases. Each contig derived from mitochondrial genomes was annotated using the MITOS webserver (Bernt *et al*., 2013). The 13 protein-coding genes, two rRNA genes, and 22 tRNA genes were aligned with MAFFT v 7.305 (Katoh *et al*., 2002) using default settings. The alignments were concatenated and the third codon position of protein-coding genes was removed. The dataset was partitioned into four subsets: one for the first codon position of protein-coding genes, one for the second codon position of protein-coding genes, one for the two rRNA genes, and one for the 22 tRNA genes. A Bayesian phylogenetic tree was generated using BEAST v 2.4.8 (Suchard *et al*., 2018). We used an uncorrelated relaxed lognormal clock model (Drummond *et al*., 2006), and a Birth Death speciation process as tree prior (Gernhard, 2008). The molecular clock was calibrated using nine fossil calibrations used by Bucek *et al*., (2019) (Table S19). The fossil calibrations were implemented as exponential priors on node times. Because transcriptome-based phylogenies unambiguously support the monophyly of Sphaerotermitinae and Macrotermitinae (Bucek *et al*., 2019) (unlike mitochondrial genome-based phylogenies; Bourguignon *et al*., 2017), we constrained Sphaerotermitinae + Macrotermitinae to be monophyletic. Similarly, we constrained non-Stylotermitidae Neoisoptera to form a monophyletic group. The MCMC chain was sampled every 1000 steps over a total of 0.4 billion generations. The convergence of the chain was assessed using Tracer v 1.7.1 (Rambaut *et al*., 2018), and the initial 10 percent was discarded. We carried out two replicate MCMC runs to ensure convergence of the chain.

### Reconstruction of Metagenome Assembled Genomes

We reconstructed Metagenome Assembled Genomes (MAGs) from metagenomes contigs using CONCOCT v 0.4.0 (Alneberg *et al*., 2014) implemented in the metawrap software v 0.9 (Uritskiy et al. 2018) with default parameters. MAG quality checking, based on 43 single-copy marker genes (Table S9), was performed with CheckM v 1.0.11 (Parks *et al*., 2015). High-quality MAGs, medium-quality MAGs, and low-quality MAGs with upward of 30% completeness and downward of 10% contamination were retained (Table S9) (Bowers *et al*., 2017). We retained low-quality MAGs that were at least 30% complete because, in some cases, they were endowed with complete pathways. Despite having fewer single-copy marker genes, 65.35% of these MAGs possessed more than 10 transfer RNA genes (tRNA) and 17.54% had at least one of the three ribosomal RNA genes (rRNA). All MAGs that did not meet these criteria were discarded. In addition, we discarded MAGs with obvious mismatches among marker genes. To identify these MAGs, we built Maximum Likelihood phylogenetic trees for all 43 single-copy marker genes with FastTree v 2.1.11 (Price *et al*., 2009). MAGs that fall in different phyla for different marker genes were considered as having obvious mismatches and were discarded. The rRNA genes were extracted using METAXA2 software (Bengtsson-Palme *et al*., 2015), tRNA genes were predicted via tRNAscan-SE tool (Chan and Lowe, 2019), and MAG coverage was calculated with the “metawrap quant_bins” command of the metawrap software (Uritskiy *et al*., 2018).

### Taxonomic annotation

The annotation of genomic features of bacterial and archaeal contigs and MAGs was carried out with Prokka v 1.14 (Seemann, 2014). This step allowed the identification of coding sequences (CDS), ribosomal RNAs (rRNA), and transfer RNAs (tRNA), which were used in downstream analyses. To identity the taxonomy of the metagenome contigs, we taxonomically annotated single-copy marker genes and other protein-coding genes in contigs longer than 1000bps. 40 single-copy marker genes were extracted using mOTU software ver1 (Sunagawa *et al*., 2013; Wu *et al*., 2013). Single-copy marker genes were taxonomically annotated using DIAMOND BLASTp (Buchfink *et al*., 2015) with e-value ≤ 1e-24 and output format 102, which uses the lowest common ancestor algorithm for annotation. Other protein-coding genes were annotated using the same settings as marker genes but with DIAMOND BLASTx algorithm. Both annotations were performed using the GTDB ver 95 database as reference (Parks *et al*., 2020). Taxonomic annotation of MAGs was based on bacterial and archaeal reference trees using GTDB-Tk v1.3.0 based on GTDB ver 95 (Chaumeil *et al*., 2020).

We used the genomic DNA extracted from whole termite guts to produce 16S rRNA gene PCR amplicon sequences. PCR reactions were carried out using the primer pairs 515F (XXXXXGTGTGYCAGCMGCCGCGGTAA, Parada *et al*., 2016) and 806R (XXXXXXXXCCGGACTACNVGGGTWTCTAAT, Apprill *et al*., 2015). All pairs of primers were endowed with unique dual tag indexes (8X overhang on the forward primer and 5X overhang on the reverse primer) to minimize the effects of index hopping between libraries. We conducted PCR amplifications using Takara Tks Gflex DNA Polymerase with the following conditions: initial denaturation (3 min at 94°C), 30 cycles of amplification (45 s at 94°C, 60 s at 50°C, and 90 s at 72°C), and a terminal extension (10 min at 72°C). All PCR reactions were scaled down using one half of the reagents recommended in the manufacturers protocol. Prepared libraries were mixed in equimolar concentration and paired-end-sequenced on the Illumina MiSeq platform. The analysis of the 16S rRNA gene amplicon sequences was performed with mothur v1.44.1 (Schloss et al., 2009) following the standard procedure for Illumina data analysis described by Kozich et al. (2013). After removing low-quality reads and chimera, sequences were clustered into operational taxonomic units (OTUs) at a sequence similarity level of 97% using VSEARCH (Rognes et al., 2016). Sequences were classified using the naïve Bayesian classifier (Wang et al., 2007) implemented in mothur and the SILVA reference database release 138 (Quast et al., 2013). The abundance of every family inferred from both 16S rRNA gene amplicon data and metagenomic data was then compared. In total 143 prokaryote lineages received identical family-level annotation in both datasets.

### Functional annotation

We carried out functional annotation of the CDSs identified with Prokka v.1.14.5 (Seemann, 2014) for all contigs and MAGs that were taxonomically annotated as bacteria or archaea using the “metagenome” option. We used the CAZy database (Lombard *et al*., 2014) as a reference to identify CDSs with carbohydrate metabolizing properties. Protein sequences were searched against a set of profile Hidden Markov models (HMM) representing CAZy domains deposited in the dbCAN database release 7 (Yin *et al*., 2012). We used an e-value lower than e-30 and coverage greater than 0.35 as thresholds to extract best domain matches.

Hydrogenases were annotated by means of HMM searches against the Pfam database version 32.0 (El-Gebali *et al*., 2019) using an e-value cut-off of e-30. Catalytic subunits of hydrogenases were classified into different classes using the k-nearest neighbor algorithm implemented in the HydDB webtool (Søndergaard *et al*., 2016). For the [FeFe] hydrogenase Group A4, we carried out manual inspection of the conserved motifs in the protein sequence (Schuchmann *et al*., 2018).

We reconstructed prokaryotic metabolic pathways from our metagenomes with KOFam scan v.1.1.0 (Kanehisa *et al*., 2016; Graham et al. 2018). We used the KEGG database as a reference and e-value cut-off of e-30. Each protein sequence was annotated to gene family level with the KEGG-Decode python module (Graham *et al*., 2018). The MAG metabolic pathways were annotated with KOFam scan v.1.1.0 using default settings. As some MAG gene families appeared to be absent after annotation against the KEGG database, to confirm, or reject, the absence of these gene families, we carried out BLAST searches (Amino acid identity >60% and alignment length > 100 amino acids) against the Annotree protein sequence database (Mendler *et al*., 2019).

### Relative abundance of gene families

The relative abundance of CDSs was calculated by mapping the raw reads on the sequences. Briefly, the reads were mapped to the assembled contigs annotated as bacteria or archaea. Relative abundance was calculated for each CDS using Salmon v.1.4.0 with the “meta” option. Salmon corrects for GC-content bias, gene-length differences, and sampling effort (Srivastava *et al*., 2020). Relative abundance of CDSs obtained as Transcripts per Million (TPM) values were retained for downstream analysis if they were embedded into contigs longer than 1000 bps and had more than 1 TPM value. Individual TPM counts were normalized using centered log(2)-ratio (clr) transformation to account for the compositional structure and unequal numbers of reads in our metagenome data. Clr transformation enhances sub-compositional comparisons (gene vs gene, bacteria vs bacteria) and reduces spurious correlations. Positive and negative TPM values indicate positive and negative departure from the overall compositional mean, which is zero (Gloor *et al*., 2017). Clr transformation of marker genes and functional genes was performed using the R package *propr* using 0.65 as a pseudo count to account for zero values (Quinn *et al*., 2017). We did not calculate TPM for MAGs, but instead used presence/absence to investigate pathway completeness.

### Statistical Analysis

We investigated whether the abundance of the genes and pathways of interest was phylogenetically autocorrelated to the time-calibrated tree of termites. To do so, we calculated the Moran’s I phylogenetic autocorrelation index using the R package *phylosignal* (Keck *et al*., 2016) on CDSs embedded in contigs longer than 1000 bps and with TPM value higher than 1. This analysis was carried out for each bacterial and archaeal phylum present in at least 5% of the metagenomes, using the combined 40 single-copy marker genes (see Table S3). A 5% false discovery rate (FDR) correction was calculated using the p.adjust function implemented in the R package *stats* (R Core Team, 2014). Similarly, we calculated the Moran’s I phylogenetic autocorrelation index for each 211 CAZymes present in more than 10% of gut metagenomes and carried out a 5% false discovery rate FDR correction. Finally, the analysis was performed for each gene involved in the reductive acetogenesis, sulfate reducing, nitrogen recycling and nitrogen fixating pathways, and the *mcrABG* gene of the methanogenesis pathway combined. We applied a 5% FDR correction.

To examine whether the abundance of the genes and pathways of interest differed with termite diet and presence of non-prokaryotic co-symbionts, we performed phylogenetic ANOVA using the procD.pgls function implemented in the R package *geomorph* (Adams and Otárola-Castillo, 2013). A 5% FDR correction was calculated using the p.adjust function implemented in the R package *stats* (R Core Team, 2014). Termite diet was determined based on literature data (Donovan et al. 2001, Bourguignon et al. 2011), and was considered as made of wood or soil. Wood-feeding termite species included feeding groups I and II (including grass and leaf litter), while soil-feeding termites included feeding groups III and IV (*sensu* Donovan et al. 2001). Non-prokaryotic co-symbionts are found in two groups of wood-feeding termites: the lower termites, which include all termites with the exclusion of Termitidae and host cellulolytic flagellates in their gut, and the Macrotermitinae, a subfamily of Termitidae that cultivates cellulolytic *Termitomyces* in fungal combs. Therefore, we recognized four groups of termites: the lower termites (LT), the soil-feeding termites (all Termitidae, SF), the Macrotermitinae (FC), and the non-Macrotermitinae wood-feeding Termitidae (WF). Similar analysis was performed on prokaryotic lineages encoding CAZyme families present in more than 10% of termite gut metagenomes in contigs longer than 5000 bps, to ensure correct taxonomic annotation. All metagenome contigs longer than 5000 bps with dinitrogen-fixing genes were also examined.

We visualized termite samples according to the abundance of CAZyme families present in their gut metagenomes using Principal Component Analysis (PCA). The PCA was performed using the prcomp function implemented in the R package *stats* (R Core Team, 2014) and visualized using the R package *ggbiplot* (Vu, 2011). Similar analyses were performed on the genes involved in reductive acetogenesis, sulfate reduction, dissimilatory nitrate reduction, urea degradation, glutamate biosynthesis, arginine biosynthesis, ammonia transport, nitrogen fixation, and *mcrABG* genes of the methanogenesis pathway.

### Uricase genes encoded by termites

We searched the 53 termite transcriptomes previously published by Buček et al. (2019) for the presence of uricases. These transcriptomes were either derived from whole worker bodies or from worker heads, and included species of all termite families. Protein sequences of predicted uricases from termites (XP_023702357, GFG34960), cockroaches (PSN45555, CDO39394), fireflies (KAF529609, XP_031344605), sawflies (XP_015591878, XP_015521616), ant (XP_011159093), fruit fly (NP_476779), and rat (NP_446220) were used as a query in TBLASTn searches. The longest open reading frames for all significant TBLASTn search hits (E-value < 10^-30^) were identified and translated using hmmer2go obtained from https://github.com/sestaton/HMMER2GO. The nonsense proteins that did not provide any significant BLASTx hit against NCBI RefSeq database (E-value < 10^-10^) were discarded. The remaining predicted protein sequences, derived from 23 transcripts, were assigned KEGG annotations using eggNOG-Mapper version 4.5 (Huerta-Cepas *et al*., 2016). The protein sequences were aligned using CLUSTAL W (Larkin *et al*., 2007) and the alignment was visually inspected.

---

**Figure S1.**
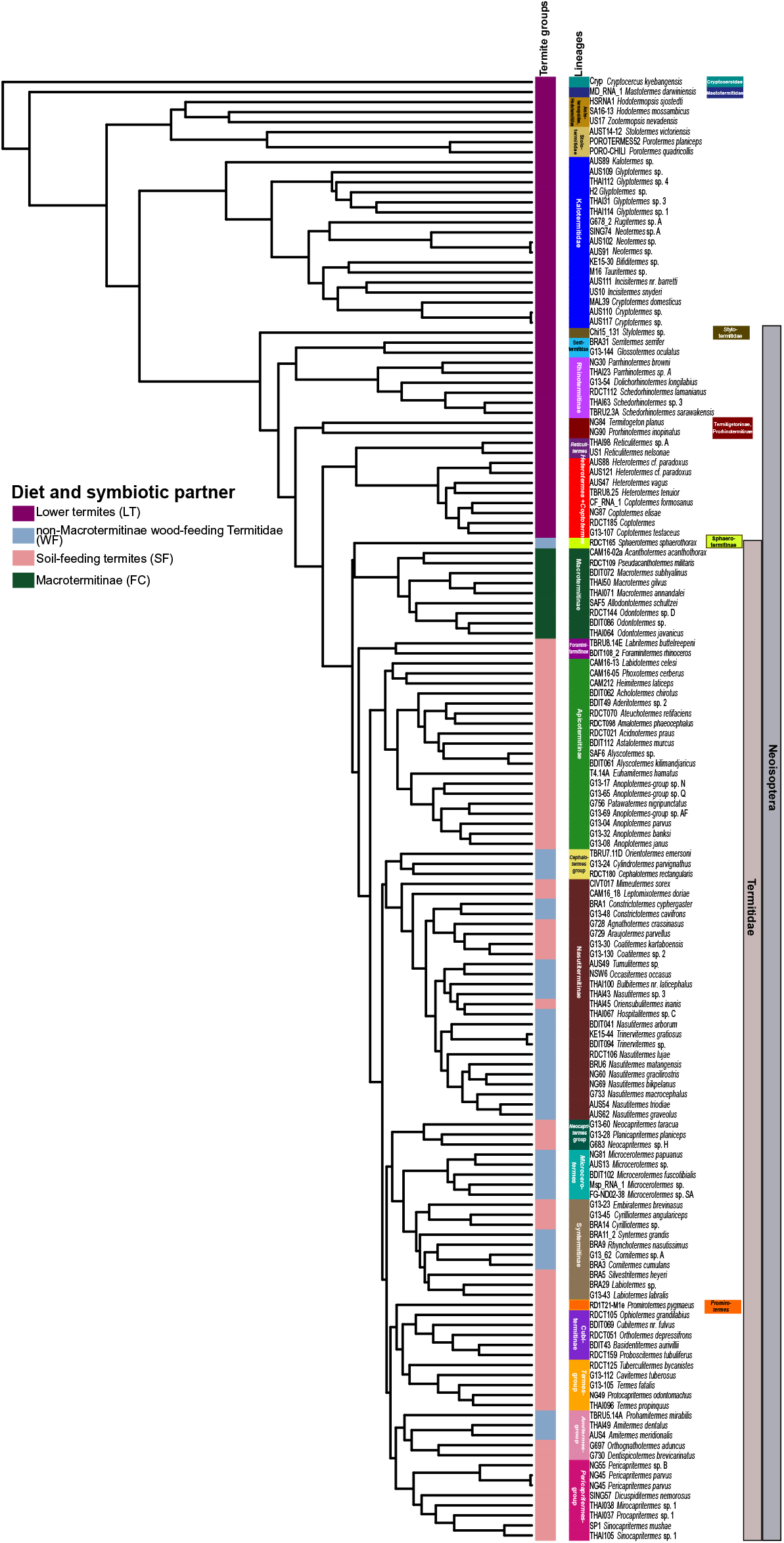
Time-calibrated phylogenetic tree of termites inferred from mitochondrial genome sequences.

**Figure S2.**
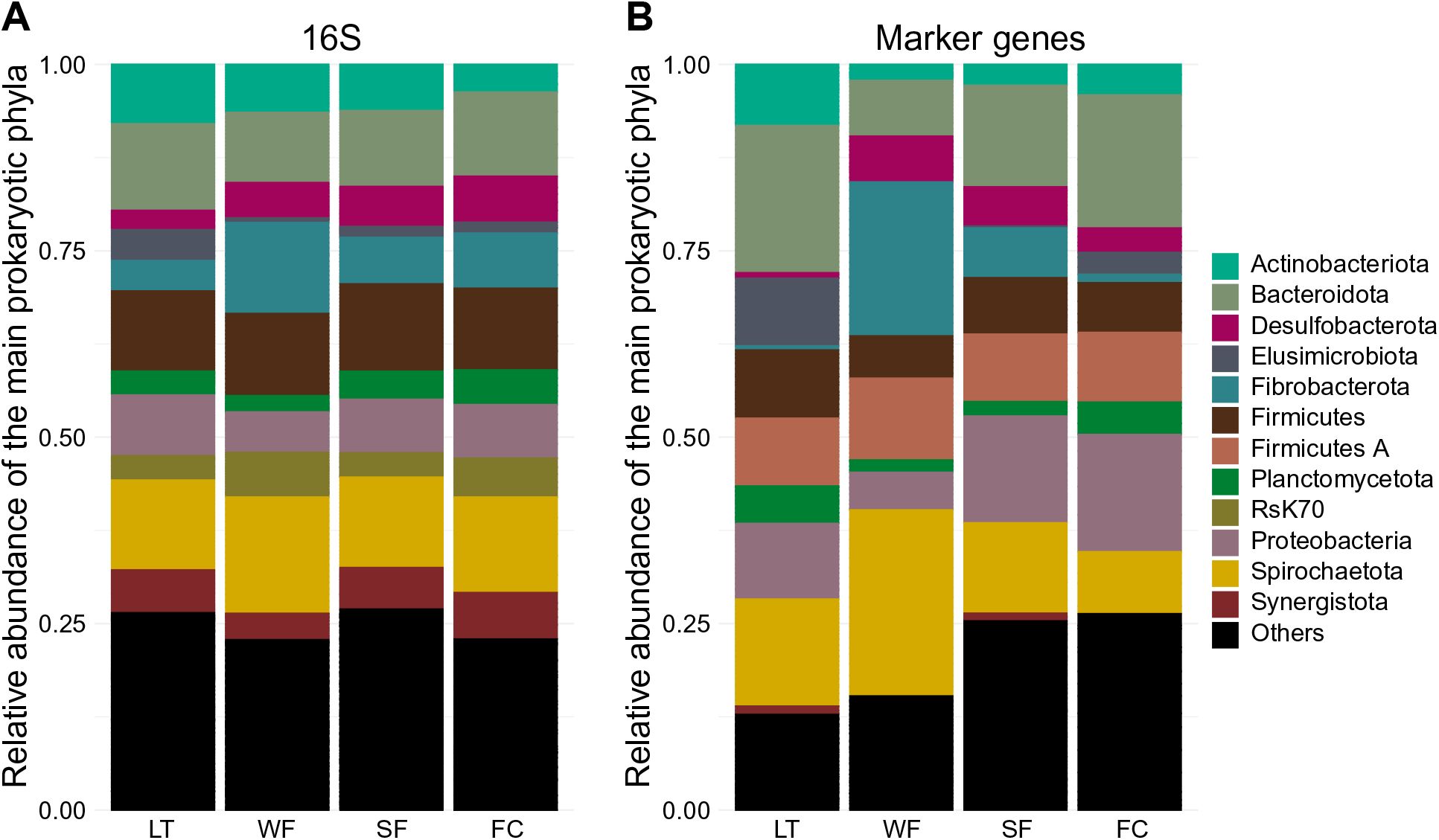
Relative abundance of archaeal and bacterial phyla inferred from the termite gut metagenomes and the 16S rRNA amplicon data of 74 termite samples.

**Figure S3.**
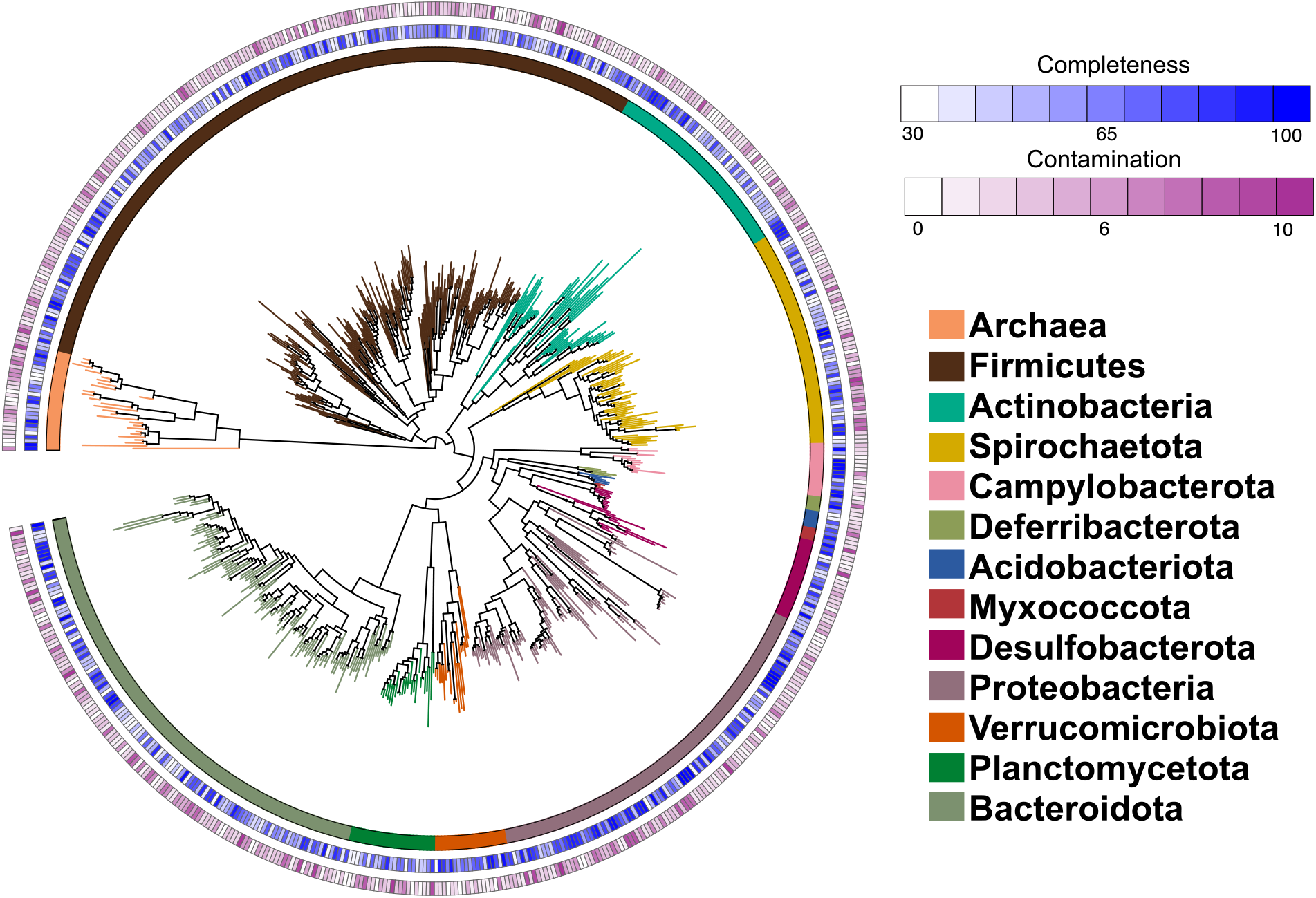
Maximum likelihood phylogenetic tree inferred from 43 single-copy marker genes of 654 metagenome-assembled genomes (MAGs). The completeness and contamination of MAGs was inferred with CheckM (Park *et al*., 2015). Detailed information about each MAG is available in Table S9.

**Figure S4.**
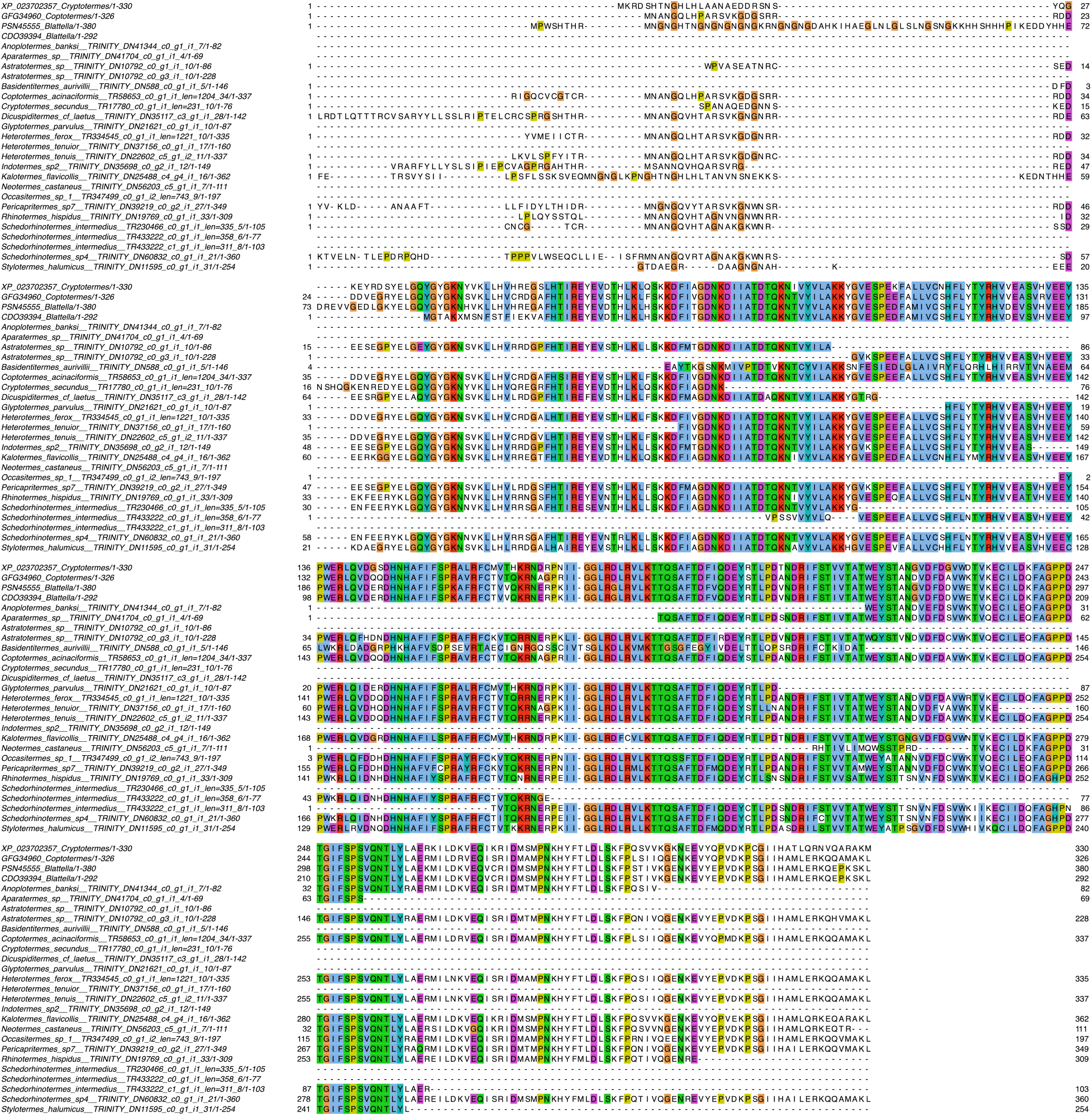
Protein sequence alignment of predicted uricases from 53 termite transcriptomes previously published in Buček et al. (2019).

**Table S1. Termite samples sequenced in the study.**

**Table S2. Relative abundance of family-level prokaryotic taxa inferred from gut metagenome and 16S rRNA amplicon data of 74 termite samples.** The prokaryotic taxonomy was determined with GTDB for marker genes and with SILVA for 16S rRNA data. The relative abundance was clr-transformed to account for differences in sequencing method and sequencing depth among metagenome samples.

**Table S3. Taxonomic distribution of major bacterial and archaeal groups based on relative abundance of 40 single-copy marker genes.** We analyzed the marker genes present in contigs longer than 1000 bps in >5% of gut metagenomes. The relative abundance is represented as transcripts per million (TPM).

**Table S4. Moran’s I phylogenetic autocorrelation index calculated for 123 prokaryote families.** Significance was assessed with 9999 random permutations. P-values <0.05 are indicated by asterisks.

**Table S5. Relative abundance of microbial CAZymes in gut metagenomes with upward of 10000 contigs longer than 1000 bps.** Relative abundance is given as transcripts per million (TPM).

**Table S6. Moran’s I phylogenetic autocorrelation index calculated for 211 prokaryotic CAZymes present in more than 10% of gut metagenomes.** Significance was assessed with 9999 random permutations. P-values <0.05 are indicated by asterisks.

**Table S7. Phylogenetic ANOVA calculated for 211 prokaryotic CAZymes present in more than 10% of gut metagenomes.** Significance was assessed with 9999 random permutations. P-values of phylogenetic ANOVA and pairwise comparisons were adjusted at 5% false discovery rate (FDR). The relative abundance of each CAZyme for the four termite groups are indicated by mean TPM values. Significance of pairwise comparisons between termite groups are indicated by asterisks (* p < 0.05; ** p < 0.01; *** p < 0.001).

**Table S8. Phylogenetic ANOVA comparing the taxonomic origin of the 19 prokaryotic CAZymes found in 10% of gut metagenomes and embedded in contigs longer than 5000 bps.** Significance was assessed with 9999 random permutations. The relative abundance of each CAZyme for the four termite groups are indicated by mean TPM values. Significance of pairwise comparisons between termite groups are indicated by asterisks (* p < 0.05; ** p < 0.01; *** p < 0.001).

**Table S9. Information about the 654 MAGs reconstructed in this study.**

**Table S10. Distribution of polysaccharide utilization loci (PULs) across the MAGs.** PULs with at least one GH and Bacteroidota PULs with at least one susCD complex are shown. MAGs containing PULs with all the components are highlighted in grey.

**Table S11. Moran’s I phylogenetic autocorrelation index and phylogenetic ANOVA performed on the genes involved in the final steps of the lignocellulose digestion in the gut of termites.** For genes composed of multiple subunits, all subunits were summed together. Significance was assessed with 9999 random permutations. P-values were adjusted at 5% false discovery rate (FDR). The relative abundance of each gene for the four termite groups are indicated by mean TPM values. Significance of pairwise comparisons between termite groups are indicated by asterisks (* p < 0.05; ** p < 0.01; *** p < 0.001).

**Table S12. Distribution of genes involved in reductive acetogenesis among MAGs.** Distribution is shown as presence (1) and absence (0). Asterisks indicate genes that were annotated using BLASTx search against the AnnoTree database (perc. identity >60%, align. length >100 aa). Other genes were annotated using HMM search against the KEGG or Pfam databases. [FeFe] hydrogenase GroupA4 were annotated using the Hyddb webtool followed by manual inspection of the conserved motifs. The total number of HycB3 (PF13247) found in each MAG is shown. MAGs with almost complete reductive acetogenesis pathway (>5 genes) and HDCR complex are highlighted in grey.

**Table S13. Relative abundance of methyl-coenzyme M reductase (*mcrABG*) gene complex present in metagenome contigs longer than 5000 bps.** Contigs were annotated using BLASTx search against the GTDB database. Relative abundance of the gene family is shown as raw TPM.

**Table S14. Distribution of genes involved in methanogenesis among MAGs.** Distribution is shown as presence (1) and absence (0). Asterisks indicate genes that were annotated using BLASTx search against the AnnoTree database (perc. identity >60%, align. length >100 aa). Other genes were annotated using HMM search against the KEGG or Pfam databases. Highlighted MAGs have a complete Methanogenesis pathway.

**Table S15. Distribution of genes involved in sulfate reducing among MAGs.** Distribution is shown as presence (1) and absence (0). Asterisks indicate genes that were annotated using BLASTx search against the AnnoTree database (perc. identity >60%, align. length >100 aa). MAGs with complete sulfate reducing pathway are highlighted.

**Table S16. Genes involved in nitrogen metabolism and fixation found in our MAGs.** Distribution is shown as presence (1) and absence (0). Asterisks indicate genes that were annotated using BLASTx search against the AnnoTree database (perc. identity >60%, align. length >100 aa). MAGs with complete nitrogen fixation or dissimilatory nitrate reduction pathways are highlighted.

**Table S17. Contigs endowed with a NifHDKENB (*nifHDKENB*, *vnfHDKENB*, or *anfHDKENB*) gene complex found in gut metagenomes.** The relative abundance is given as raw TPM.

**Table S18. Contigs endowed with a NifHDK (*nifHDK*, *vnfHDK*, or *anfHDK*) gene complex found in termite gut metagenomes.** The relative abundance is given as raw TPM.

**Table S19. Fossil calibrations used to calibrate the time-calibrated tree of termites.**

